# From Diversity to Complexity: Microbial Networks in Soils

**DOI:** 10.1101/2021.12.14.472586

**Authors:** Ksenia Guseva, Sean Darcy, Eva Simon, Lauren V. Alteio, Alicia Montesinos-Navarro, Christina Kaiser

**Affiliations:** Centre for Microbiology and Environmental Systems Science, University of Vienna, Vienna, Austria; Centro de Investigaciones sobre Desertificación (CIDE, CSIC-UV-GV), Carretera de Moncada-Náquera Km 4.5 46113 Moncada, Valencia, Spain

**Author notes:** Corresponding author (K. Guseva); (C. Kaiser).

**Keywords:** Microbial network analysis, Co-occurrence networks, Ecological networks, Microbial community structure, Soil microbial ecology

## Abstract

Network analysis has been used for many years in ecological research to analyze organismal associations, for example in food webs, plant-plant or plant-animal interactions. Although network analysis is widely applied in microbial ecology, only recently has it entered the realms of soil microbial ecology, shown by a rapid rise in studies applying co-occurrence analysis to soil microbial communities. While this application offers great potential for deeper insights into the ecological structure of soil microbial ecosystems, it also brings new challenges related to the specific characteristics of soil datasets and the type of ecological questions that can be addressed. In this Perspectives Paper we assess the challenges of applying network analysis to soil microbial ecology due to the small-scale heterogeneity of the soil environment and the nature of soil microbial datasets. We review the different approaches of network construction that are commonly applied to soil microbial datasets and discuss their features and limitations. Using a test dataset of microbial communities from two depths of a forest soil, we demonstrate how different experimental designs and network constructing algorithms affect the structure of the resulting networks, and how this in turn may influence ecological conclusions. We will also reveal how assumptions of the construction method, methods of preparing the dataset, and definitions of thresholds affect the network structure. Finally, we discuss the particular questions in soil microbial ecology that can be approached by analyzing and interpreting specific network properties. Targeting these network properties in a meaningful way will allow applying this technique not in merely descriptive, but in hypothesis-driven research.

## 1. Introduction

Networks present a powerful mathematical framework to explore the complexity of relationships within an ecological community. Their potential lies in a formal but intuitive representation of the organization of complex systems, where entities (f.e. species) are displayed as nodes, and interactions or associations among them as edges (links). Since the beginning of the century, network analysis has become an established tool for investigating species relationships in ecosystems (MAY (1974); Bascompte (2009); Poisot et al. (2016)) and has experienced an upswing in soil ecological research over the past ten years (Fig. S1).

One of the original aims of the field is to reveal interplay between species interactions present in nature, which allows to gain valuable information on ecosystem properties. Data for ecological network analysis of species interactions is ideally obtained by direct observation of physical contact, f.e. pollination of plants by insects or birds, or mycorrhizal fungi colonizing plant roots. However even in such cases identifying species interactions within an ecosystem can be challenging, as it requires repeated observations or a mechanistic understanding of them. An alternative approach for investigating networks of species relationships has been the analysis of spatial co-occurrence patterns of species with so-called co-occurrence networks. Although the use of co-occurrence data often limits possible conclusions on species interactions, it contains valuable information regarding community structure and assembly mechanisms which can be explored by the use of network analysis (Blanchet et al. (2020)).

In microbial ecosystems, associations between microbial taxa can usually only be assessed via co-occurrence data, as interactions between microbes in their natural environment are much more difficult to observe than in macro-ecosystems. Their assessment would require complicated experiments, for example, involving records of events of feeding or cross-feeding, mechanisms behind biofilm formation or other types of interactions. All of which are impossible to carry out in situ, particularly in complex environments such as the soil. Network analysis in microbial ecology therefore needs to rely on the observation of species co-occurrence or co-exclusion patterns within molecular microbial datasets (f.e. derived from 16S amplicon sequencing) across a sufficiently high number of environmental samples (Röttjers and Faust (2018)). Networks generated from this indirect information are known as ‘microbial co-occurrence networks’ or ‘microbial association networks’, and have gained much popularity in microbial ecology in the last decade (Fig. S1). Analogous to their macro-ecological counterparts, the observed structure of microbial co-occurrences is the result of community assembly, a complex ecological mechanism, which is influenced by various processes such as environmental filtering (f.e. species that respond in the same way to an environmental factor tend to co-occur in samples with variation of that environmental factor), species interactions (f.e. two microbial taxa need to exchange specific metabolites for increasing their fitness), dispersal dynamics and stochastic processes (Faust (2021)). Network analysis of in situ distribution of microbial taxa in soil reveals the final result of these community assembly processes, and can therefore be a great starting point to better understand them. To interpret these networks it is important to distinguish microbial associations from microbial interactions (Barner et al. (2018)). While the first refers to the observed statistical signal obtained from co-occurrence (or abundance) patterns, microbial interactions constitute the relationships between species present in natural systems.

The advent of high-throughput sequencing techniques has led to widespread use of co-occurrence analysis of microbial communities, which went along with the development of significant methodological advances. It is currently used in a rapidly growing number of soil microbial ecology studies. While not more than 10% of microbial network analysis studies were conducted on soil datasets before 2010, this number has steadily increased since then to 35% in 2020, see (Fig. S1) of Supplementary Material. While this approach has great potential to enrich soil studies, the intrinsic complexity of soil adds new challenges to the construction and interpretation of network models. The difficulty consists in taking into account the inter- and intra-variability of samples, which are a result of soil heterogeneity (Carr et al. (2019)). The first leads to a high number of environmentally driven association patterns, which can be mistaken as species interactions ((Armitage and Jones (2019)). The second, refers to the use of large sample volumes, where naturally segregated and chemically diverse micro-habitats are mixed (usually 0.25 cm^3^ soil for DNA analysis, derived from a homogenized sampling volume of 250-500 cm^3^ soil). The resulting loss of information about the small-scale physical, chemical and biological diversity of soil may also confound the signal of biological interactions (Berry and Widder (2014)).

Overall, network analysis relies on a clear understanding of what the constructed network model represents and the data used for it (Poisot et al. (2016)). Unfortunately, due to misuse, networks have unfairly acquired a mixed reputation in soil microbial ecology. On one hand they are considered essential for understanding the structure and properties of microbial communities (Faust and Raes (2012)). On the other hand, theoretical ecologists warn against inappropriate conclusions derived from inaccurate data handling, unsuitable inference methods and more often an inappropriate interpretation of the obtained network model (Carr et al. (2019); Berry and Widder (2014); Blanchet et al. (2020)). As the use of networks in studies of soil microbial communities continues to increase, we see an urgency to discuss the challenges the approach inherits from other fields and new challenges the soil system poses.

In this Perspective we describe the main ideas behind network inference from soil microbial co-occurrence datasets (Sec. 2), the network construction itself (Sec. 3), and the mathematical framework for the analysis of such networks (Sec. 4). After this overview, we discuss the challenges of application of these methods to soil microbial communities (Sec. 5). The network construction is illustrated using a dataset of microbial communities from different depths of a forest soil. We discuss how differences in the experimental design, data preparation and filtering, as well as network construction algorithms affect the structure of the resulting networks. We also discuss how these differences affect our interpretation of the edges in the network and influence our ecological conclusions. We argue that to exploit the full potential of network analysis for soil ecological studies it takes a combined effort from both experimental and theoretical sides. Applying network properties in a meaningful way enables the application of network analysis not only in merely descriptive, but also in hypothesis-driven research.

## 2. Networks as models of microbial communities in soil

Microbial association networks are constructed based on co-occurrences of microbial taxa across a sufficiently high number of environmental samples (BOX 1). Each node in the constructed network represents one microbial taxonomic unit while an edge between two nodes represents a significant association between these microbial taxa. There are different ways how network construction algorithms determine whether two taxa are significantly associated, and thus connected by an edge in the network. The most common approach is based on pairwise correlations among taxa across all samples. (for details see Sec. 3 and BOX 1). One of the key prerequisites for interpreting the structure of the resulting network is to understand what an edge, i.e. a significant association, represents.

### BOX 1

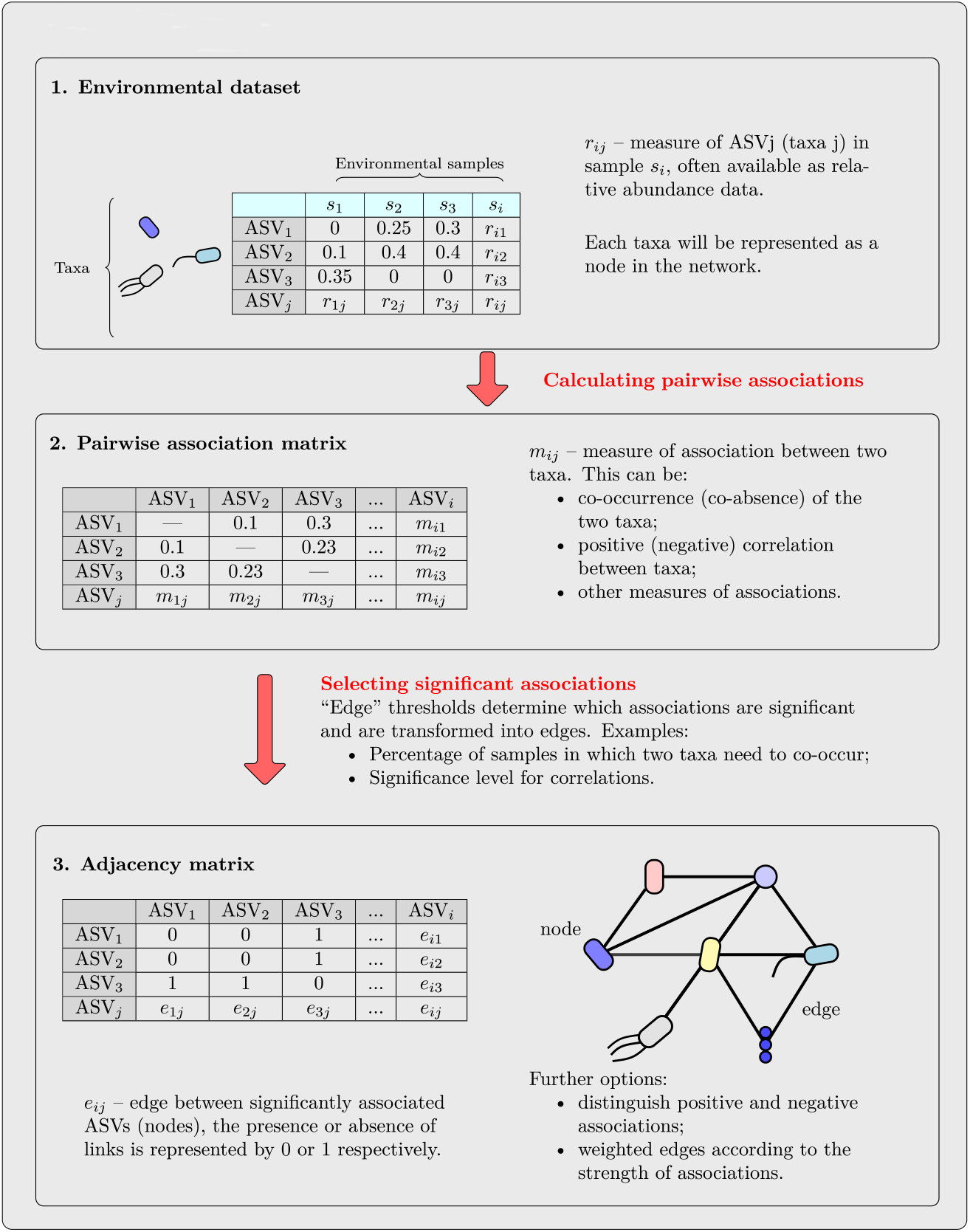

To understand what an edge (observed association) could potentially mean we need to understand the mechanisms that lead to the observed co-occurrence pattern, i.e. the mechanisms by which microbial communities assemble. Microbial community assembly is driven by multiple processes, such as environmental filtering, inter-specific interactions, reproduction/mortality events, dispersal limitation and sporadic mutations (Nemergut et al. (2013)). Besides, other factors such as legacy effects, the importance of the order by which species are introduced into a system, show how community composition can depend on elusive factors (Fukami (2015)).

The combined outcome of all these hidden ecological processes leads to the spatial distribution of microbial taxa in soil. The relevance of each process for co-occurrence patterns at a given scale, however, remains controversial. Soil microbial network analysis faces the challenge to use observations of microbial distributions which are the result of multiple processes for constructing reliable models of microbial communities as schematically depicted in (Fig. 1). The process of reconstructing an underlying ecological reality from observed data by means of an abstract mathematical model is called inference. Network inference, which is also often termed network reconstruction, correspondingly looks for mathematical models that represent relationships in a system as a network with the aim to reconstruct the hidden ecological relationships. Despite the importance of building networks on a suitable mathematical model, many state-of-the-art network reconstruction methods don’t go beyond a mere identification of patterns, such as significant correlations, and thereby only provide a naive representation of the hidden reality without a clear interpretation. This is often the case for microbial co-occurrence networks in soil which are most often constructed based on pairwise correlations. To understand what an edge, that connects two taxa in a network, represents, we need to understand why these two taxa significantly co-occur across a high number of samples. Despite the wealth of underlying causes for the spatial distribution of microbial taxa in soil, significant species co-occurrences, which are the basis for network edges, are often interpreted to be caused mainly by interactions between the taxa. To avoid such misleading conclusions from network analyses, we suggest that thinking about the underlying biological processes and statistical assumptions should be at the core of network model construction of soil microbial communities. Nevertheless, we concede that the detection of associations can be a valuable intermediate step, which can help to generate hypotheses about a given system and its taxa. Finally, for a safe interpretation, it is essential to (i) consider the details of the experimental design, as well as the assumptions of the used statistical methods, as both may influence the potential meanings of network edges, and (ii) (ideally) develop network reconstruction methods that go towards a more accurate representation of the underlying ecological relationships (see Sec. 5)

**Figure 1:**
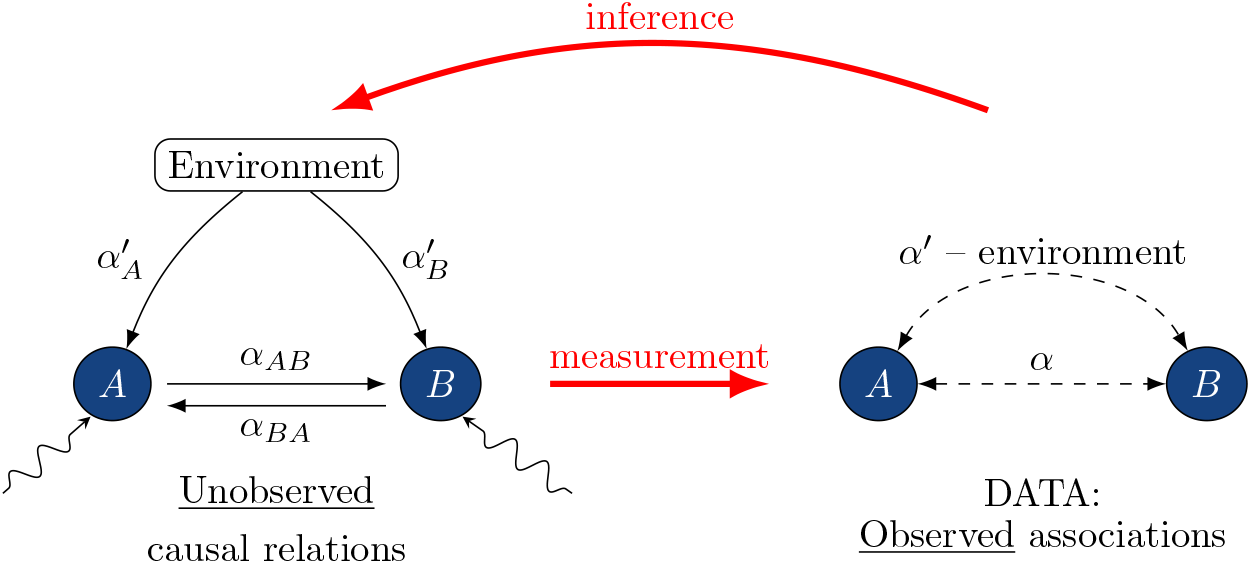
The hidden relationships (left diagram) within an ecosystem are reconstructed (inferred) from limited observations and measurements, in this case of associations between microorganisms (right diagram).

## 3. Network construction

Network model construction aims to reconstruct a network of ecological relationships existing in nature from observational data. This aim is inherently challenging and can only result in a model that in the best case reflects some aspects of the existing ecological network. While there is an agreement that a graphical representation of ecological relationships in a system is useful, there is no universal framework to infer these relationships from environmental datasets. In this section we describe the most popular statistical approaches used to assess associations between microbes and discuss their potential and drawbacks.

We illustrate the process of network construction and the effect of considering different experimental designs on the network structure by examining the microbial communities from forest soil at two depths, based on 18 samples taken from a lower soil layer (15–20 cm depth) and 20 samples from the upper soil layer (0–5 cm depth). The community composition was evaluated by sequencing 16S SSU rRNA marker gene. The description of sampling, DNA extraction and sequencing processes are given in the Supplementary material. The purpose of this analysis is to elucidate what effect i) the inclusion of a strong environmental factor which may influence the spatial distribution of microbial taxa in the soil (soil depth) and ii) the choice of a certain statistical network inference method can have on the resulting network structure and interpretation.

### 3.1. Preparation of the dataset

In most network construction approaches, associations between microorganisms are identified either from probabilities of co-presence (co-absence) or from pairwise correlations of species abundance in a number of environmental samples. Abundances of microbial taxa in natural microbial communities are usually assessed by amplicon sequencing of highly preserved, phylogenetically informative marker genes, such as the 16S ribosomal RNA for bacteria and the internal transcribed spacer (ITS) for fungi. The sequenced reads are grouped by sequence similarity into operational taxonomic units (OTUs) or recovered from sequenced reads as amplicon sequence variants (ASVs) (Callahan et al. (2017)), which can then be classified into taxonomic categories.

Two main characteristics of amplicon datasets are: (i) they are compositional, which means that abundances of taxa can only be interpreted in relative terms (Gloor et al. (2016); Morton et al. (2019), and, especially for soil microbial communities, (ii) taxa are sparsely distributed, which means that datasets contain a high amount ASVs or OTUs that only occur in a fraction of the samples (Alteio et al. (2021)). Both of these characteristics pose a major challenge to network construction approaches, as correlations carried out on relative abundance and sparsely distributed datasets can lead to spurious results (Morton et al. (2019); Alteio et al. (2021)). In the next subsections we discuss how to reduce the potential bias introduced by these features of the dataset, by applying appropriate data filtering and transforming steps before the actual network construction (Fig. 2).

**Figure 2:**
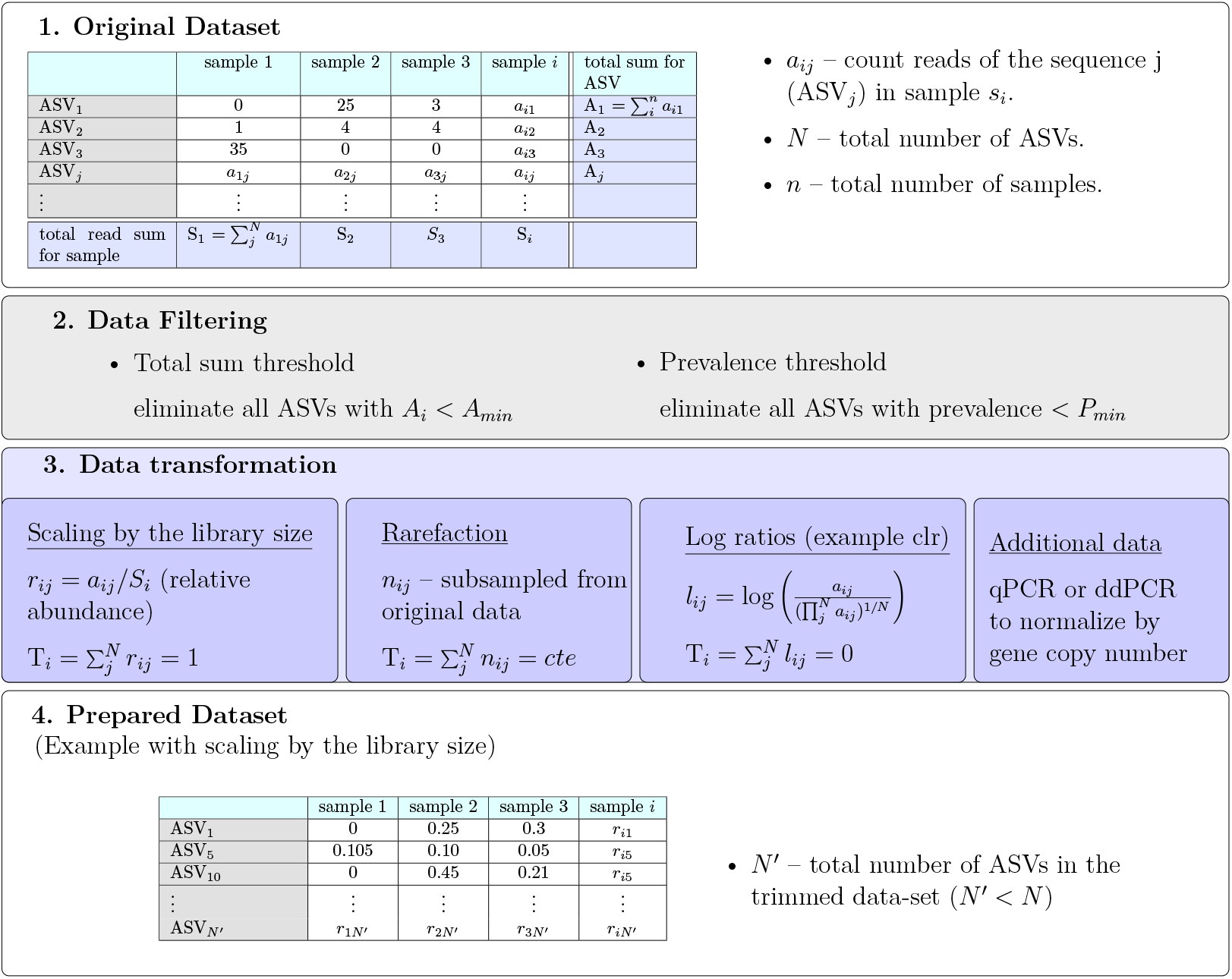
Workflow for the preparation of a dataset which precedes the network construction. It involves filtering out some ASVs and a data transformation, such as scaling the number of reads *a*_*ij*_ by the total read sum in a sample *S*_*i*_, or taking a centered log ratio (clr). The transformed dataset contains relative abundance values constrained by a constant (cte) sum Ti, or even inferred absolute abundance values for each ASV in each individual sample.

#### 3.1.1. Interpreting reads: estimations of abundances of microorganisms from high-throughput sequencing data

Amplicon sequencing data is compositional, which means that it contains information only on relative abundances. There are several methods to extract this relative information out of measured count reads, which is most often done by: (1) rescaling the count reads by the library size (*a*_*ij*_/*S*_*i*_) or (2) rarefaction, see (Fig. 2). The constant sum constraint ties all values together and consequently the obtained values in each sample are not independent. The computation of correlations on these non independent values is unreliable and thus gives rise to spurious correlations (Gloor et al. (2016); Lovell et al. (2015)). The introduced bias is stronger if the reads of some taxa dominate the data, while it plays a minor role for data sets of highly diverse communities. Constructing correlation-based networks from compositional datasets is common practice, and particularly often seen in studies of soil microbial communities. It is important to keep in mind that, depending on the dataset, these networks may be dominated by spurious associations. To overcome the problem of spurious correlations caused by compositional data, log ratios instead of transformations (1 and 2) were proposed (Fig. 2). However, they require an adequate exchange of zeros by pseudo-counts, which implicitly assumes that all unfiltered taxa are present in all samples, but were not detected due to shortcomings of sequencing.

As it is not possible to directly recover absolute abundances from sequencing data, complementing amplicon sequencing data with additional data can improve quantitative insights into microbial communities. One possibility is to use qPCR or droplet digital PCR (ddPCR) to quantify the number of marker gene copies in the sample (Perisin et al. (2016); Jian et al. (2020)). Another option is to infer marker gene copy numbers based on the phylogeny of organisms (Kembel et al. (2012)). However this step relies heavily on a good classification of the taxonomy and while the approach generally works well for gut bacteria, it can be more challenging for fungi Lavrinienko et al. (2021).

#### 3.1.2. Interpreting zeros: sparsity in amplicon sequencing datasets

Another important aspect of amplicon sequencing data sets is their sparsity, which results from biological complexity as well as technical challenges associated with nucleic acid extraction and sequencing. Particularly in soil, which is characterized by high species diversity and heterogeneity, most detected taxa are rare and only appear in a small fraction of the total number of samples. Consequently, these datasets are often highly zero-inflated, for example for our datasets close to 50% of all read values across samples are zeros. Observed zeros result from several very different processes (Kaul et al. (2017)). On one hand, there are biological zeros that show the real absence of a microorganism in the community and result from a complex interplay of community assembly and dynamics, see Sec. 2. On the other hand, the data can display technical zeros that result from upstream sample processing, or limitations of sequencing data. Upstream challenges are related to DNA extraction bias and amplicon primer design, which may not capture particular subsets of organisms. In addition, zeros can be caused by undersampling of a soil sample. As a result of the high diversity of soil, high sequencing depth must be achieved to capture the scope of diversity. The sparsity of the data and the low number of sequenced samples typical for most experiments make it difficult to establish meaningful associations among species with low prevalence. First of all, the appearance of taxa in only a few samples makes it statistically impossible to spot any negative associations among them (Cougoul et al. (2019)). Second, the observation of co-absence of taxa in samples can result in spurious positive associations. False positive associations may arise since correlations, such as the widely used Spearman’s rank, can overestimate obtained values due to many repeated zero values (tied ranks) (Connor et al. (2017)). Since we do not know how to correctly interpret zeros in our data, we should not attribute significance to these correlation values inflated by species absences. Therefore, it is common to establish a prevalence threshold, requiring e.g. a presence in 20% of all samples, or set a total sum threshold requiring a minimum number of reads, see (Tab. 1). The arbitrariness of thresholds was pointed out in (Cougoul et al. (2019)), and remains controversial in sequencing data analysis. Alternatively randomly chosen pseudocounts can be added to the dataset(Connor et al. (2017)). For our dataset of forest soil we have used a total sum threshold of 100 reads. Due to the structure of our dataset, in which taxa with low reads also were of low prevalence, this criteria eliminated the taxa of low read numbers, as well as low prevalence, leaving 595 ASVs for the combined dataset of upper and lower soil layers, as well as 283 and 304 ASVs for the separate analysis of upper and lower soil, respectively. We have checked that zeros do not have influence for the established pairwise associations in this trimmed dataset, see details in the Supplementary material.

### 3.2. Establishing associations

#### 3.2.1. Choice of metric to measure associations and choice of an edge threshold

The most common methods of network construction rely on establishing edges between taxa with significant pairwise associations. In this case the network depends strongly on two key decisions: (1) the choice of a metric to quantify the strength of association’s signal; and (2) the choice of a significance threshold – referred to as edge threshold (see BOX1), which serves as a criterium to differentiate a true signal from noise. The strength of associations (1) is most often measured using parametric Pearson or non-parametric Spearman or Kendall-τ correlation coefficients. The latter two are more convenient since they can be used on non-normally distributed datasets, as is the case of microbiomes. However these are not the only possibilities for (1) and measures such as Bray Curtis, Kullback Leibler dissimilarities or even mutual/maximal information can also be used to quantify associations. After the metric is chosen the next step is (2) — the choice of an edge threshold, τ. Before this threshold is applied, the networks are fully connected, with each node (taxa) connected to all other nodes. In these networks the weight of each edge is given by the corresponding pairwise correlation (association) value *ρ*. To illustrate this, we provide the histogram of Spearman correlation coefficients for our dataset of the upper soil core, which shows that the weights of most of such edges are close to zero and cannot be interpreted as a strong signal of an association (Fig. 3a). The solution therefore, is to eliminate edges with weights below a given edge significance threshold τ. In this way, the threshold is related to the sparsity of the network, where lower values result in very densely connected networks, while high values in networks with fewer edges, see (Fig. 3b). The choice of the threshold value can be either totally arbitrary using a single value for all edges, or it can be evaluated for each edge by comparisons with null models. In the second case, the null model is obtained by shuffling read values among taxa and samples. In other words, a new data set is obtained by randomly selecting values from the original one. The shuffling can be followed by a possible rescaling, due to new total read values in each sample. This procedure is done to break any non-random pairwise association patterns while keeping some of the original characteristics of the data (e.g. sparsity, compositionality and marginal count distributions of taxa). Each pairwise correlation value in the original data set is then ranked against the correlation value of the same pair in the shuffled dataset (Faust and Raes (2016)). This allows one to check the probability of observing such correlation by mere chance (i.e. in similar random data). The p-value determines if the edge should be kept or eliminated. (Fig. 3a) marks with light blue the pairwise correlation values which are left after such a procedure (using 1000 shuffled datasets), and demonstrates that the procedure also works as a sort of edge threshold. Another popular method for edge threshold selection is using ideas from random matrix theory (RMT, mathematical research area that studies matrices with random elements) (Deng et al. (2012)). The approach determines at which threshold the generated adjacency matrix no longer has properties compatible to that of a fully random network and uses that as edge threshold. Although the procedure takes away the arbitrariness of the choice, it also selects a hard threshold, and relies on the assumption that biological networks have a completely random structure at the low τ limit.

**Figure 3:**
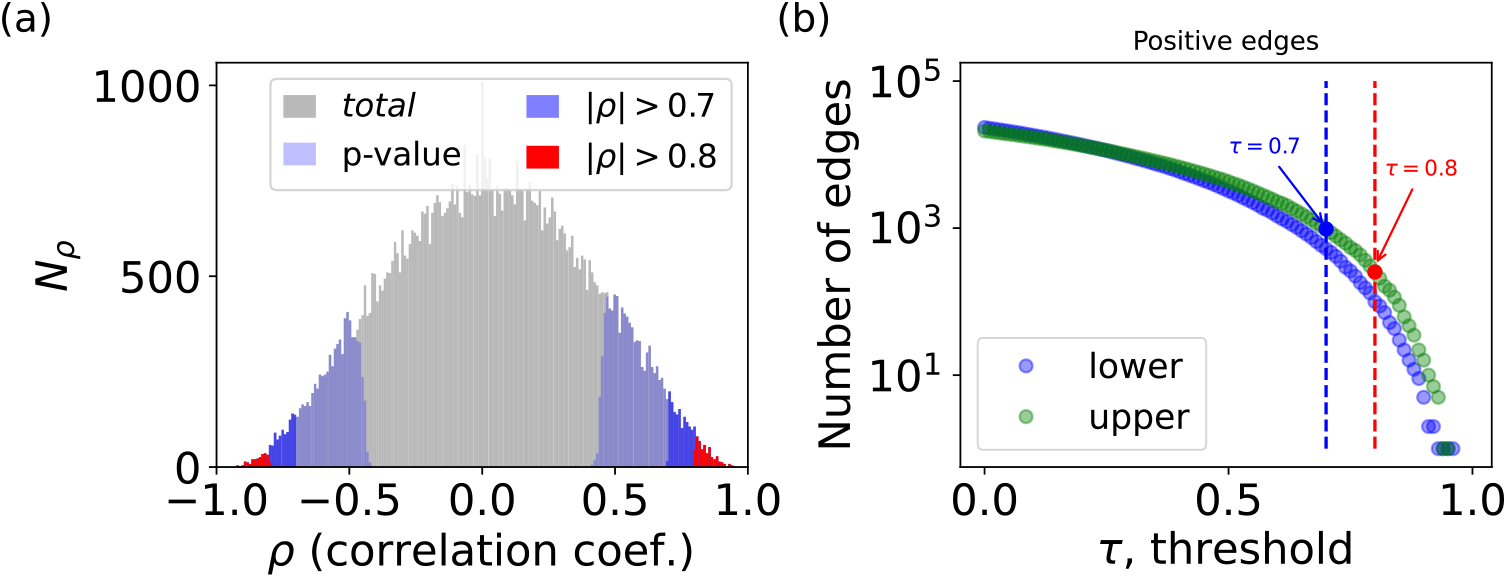
The choice of the threshold, to establish an association edge, changes the sparsity of the constructed networks. Frequency, *N*_*ρ*_ of the Spearman’s rank correlation coefficients *ρ* obtained for all ASV pairs in the upper soil core (in gray). The selected *ρ* values by the use of different edge thresholds τ are marked with different colors (red and dark blue). In light blue we show *ρ* selected as significant by comparison with a null model (for that we use 1000 shuffled versions of our dataset, and p-value = 0.05). (b) Number of positive edges (*p >* 0) in networks constructed with different edge thresholds.

We illustrate the networks obtained by the use of two thresholds τ = 0.7 and 0.8 in (Fig. 4a, c, e) and (Fig. 4b, d, f) respectively. For both thresholds, networks from upper and lower soil layers, as well as from both soil layers combined present two highly interconnected sets of nodes (modules or clusters): While the two sets are completely disconnected for τ = 0.8, we can detect candidate taxa that connect the two potential communities for the lower threshold. The choice of threshold also changes important network properties, such as mean degree, mean distance and modularity. This demonstrates that the choice of edge thresholds can influence the interpretation of the network structure in terms of its connectivity and the relevance of individual taxa in the system.

**Figure 4:**
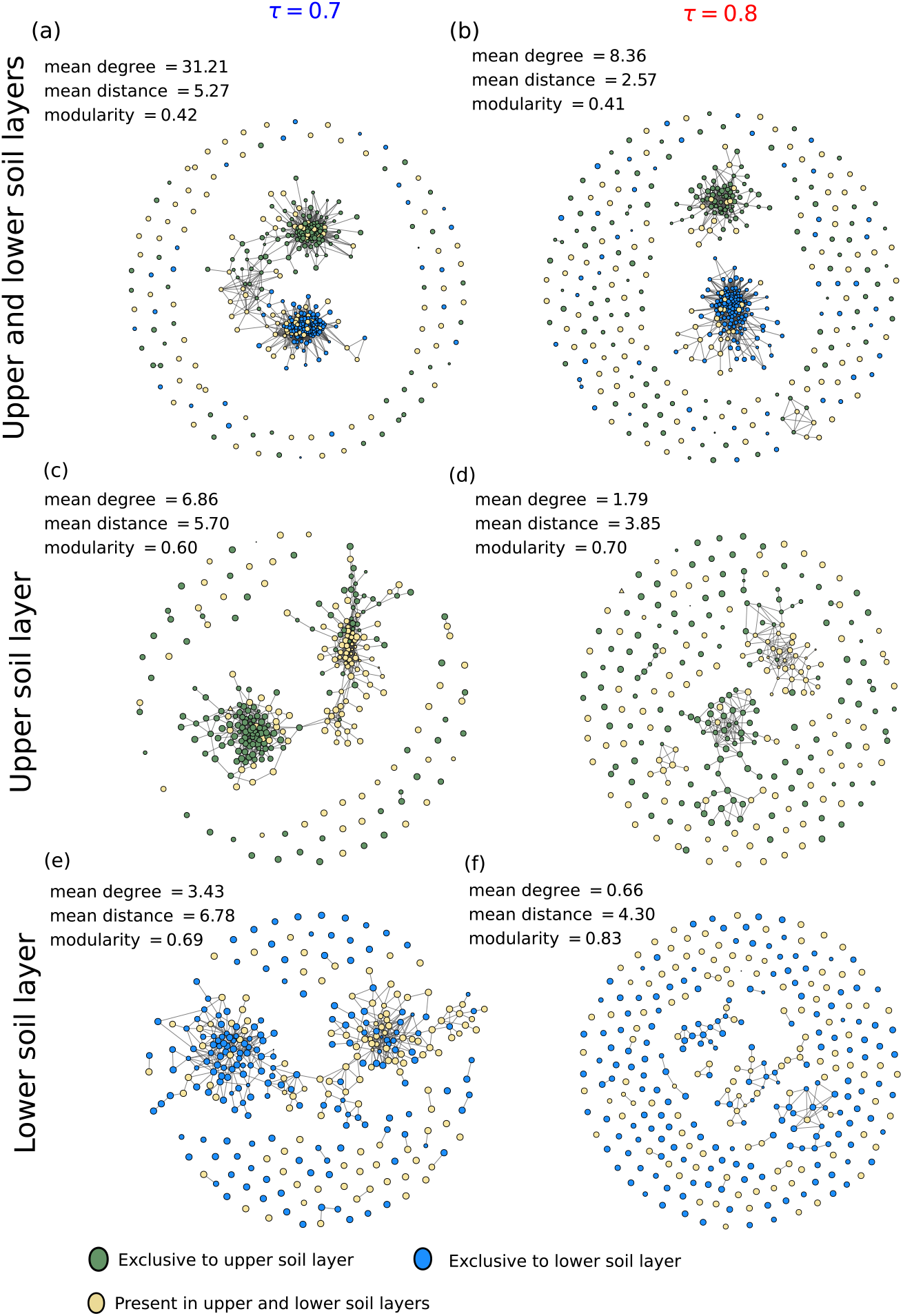
Association networks of microbial taxa in a Beech forest soil at different soil depths constructed from Spearman’s rank correlation coefficient at different edge thresholds. Shown are networks constructed from upper and lower soil layers combined (a,b), and from each soil layer separately (c-f), for edge thresholds τ = 0.7 (a,c,e) and 0.8 (b, d, f). Notice how the selected value for the edge threshold changes the number of edges in the network as well as the network structure (for details on preparation of the data set see Supplemental material). The network of the combined dataset (a, b) shows not the total 595, but only 461 ASVs shared with either one of the two other datasets (lower panels). The network with the phylogenetic classifications of nodes is also provided in the Supplement. Network visualization was done with graph-tool (Peixoto (2014)).

The advantage of using the strategies described so far is that they involve simple computations. However, as described before they suffer from several limitations: they do not take into account the compositional nature of the dataset and they cannot clearly separate association signals from noise. We can see that the choice of a threshold constitutes a bargain between the detection of spurious links (false positives) and neglecting existent associations (false negatives). The problem of threshold selection is not unique to species co-occurrence networks, but also exist in other areas from inference of gene co-expression to social networks (Couto et al. (2017)). The disadvantage of any hard threshold, independent of the selection criteria, is that the thresholding procedure itself may distort the underlying data, strongly affecting the observed network patterns (Cantwell et al. (2020)). In particular, as we explain in the next section, this occurs because there is no possible threshold that can distinguish direct from indirect associations. For microbial networks what we often see is that thresholding leads to very dense network structures (‘hairballs’ (Faust (2021))). All these reasons make it difficult to derive an ecological interpretation from the pattern in the constructed networks. Several network construction tools were developed to tackle the listed issues, we describe some of them in the next section.

#### 3.2.2. Statistical methods that improve reconstruction of ecological networks

In the last decade several ready-to-use tools were made available for network construction. Here we highlight: CoNet, which combines several association measures described in Sec. 3.2.1 (Faust and Raes (2016)); and MENA, which uses random matrix theory to decide on the threshold choice (Zhou et al.; Deng et al. (2012)); both are widely used for network construction for microbial soil communities. Another tool which stands out is SparCC due to its ability to treat possible compositionality bias in the data (Friedman and Alm (2012)). It uses log-ratio variance (vlr) to estimate associations, a transformation that presents advantages as it is subcompositionaly coherent (see definition in (Tab. 1)). More precisely this transformation allows to consistently analyze any subset of the data, without any contradictions in the results. The SparCC algorithm works under the assumption of few associations between taxa. The final adjacency matrix is evaluated again either by establishing an edge threshold or a comparison with null models.

To improve our ecological interpretation, it is also of interest to distinguish direct from indirect dependencies between taxa. For example, it is well known that two taxa may show a significant correlation because both are under the influence of a third taxa, although they do not directly interact with each other. The difficulty is that no choice of edge threshold for correlations is capable of distinguishing between direct and indirect associations. To solve this issue a new set of algorithms for network construction was proposed based on probabilistic graphical models (Kurtz et al. (2015); Fang et al. (2017); Yoon et al. (2019); Jiang et al. (2021); Yang et al. (2017)). These models represent complex probabilistic relations in the form of diagrams (graphs), where nodes represent taxa and edges represent conditional dependencies between them. Such a graph can be easily constructed for Gaussian multivariate data and for this case it is known as Gaussain graphical model (other names such as partial correlation network or concentration graph are also used). The conditional dependencies are evaluated from non-zero elements of the inverse covariance matrix (Bishop (2007)). It is important to note that strictly speaking for non-Gaussian settings, as is the case for microbial data sets, the relationship between the inverse covariance and conditional dependencies is not known. However, the approach is still used as it is expected to be informative beyond its standard domain (Liu et al. (2009)). The inverse covariance is computed with a popular statistical method known as glasso, lasso regularized maximum likelihood estimator. In this case, the sparsity of the network is tuned with a regularization parameter, λ, with small values of, λ corresponding to densely connected networks and large values to fully disconnected networks. The method in general selects a range of, λ values and generates several networks. The next step is to select the best model, considering its complexity and data fitting ability. The most well known model selection algorithms are BIC, EBIC, Stars. SPIEC-EASI (Kurtz et al. (2015)) was one of the first tool packages based on graphical models and therefore is the most well-known. However it is not the only package based on a graphical model, there is a series of other recently developed tools among them: gCoda (Fang et al. (2017)), SPRING (Yoon et al. (2019)) and BC-GLASSO (Jiang et al. (2021)). We also illustrate the outcome of network construction using SPARCC and a SPIECEASI-like (clr transform, glasso and Stars) algorithm in (Fig. 5). For simplicity, we do not show nodes which have zero edges in this image. Again, as in (Fig. 4) it is possible to identify the formation of two clusters, one with the taxa unique to the respective core depth (nodes shown in blue and green), and another with taxa shared by the two cores (nodes shown in red). Despite the differences between methods, they capture similar patterns, although only half of the edges are shared among them, see (Fig. 6). Nevertheless, the ecological processes behind the observed structure remain unclear. Note that these methods perform rather poorly when used on artificial data sets produced even from simple models of population dynamics such as gLV (generalized Lotka-Voltera) when different environmental conditions are used (Berry and Widder (2014); Hirano and Takemoto (2019)). One of the reasons is that indirect dependencies between taxa can also appear due to environmental factors, for example due to niche overlap. It is possible to introduce environmental parameters as additional nodes in the network (Faust (2021)). Another form to deal with environmental confounders is to use a model based approach as done by mLDM (Yang et al. (2017)), a tool introduced recently based on an hierarchical Bayesian statistical framework.

**Figure 5:**
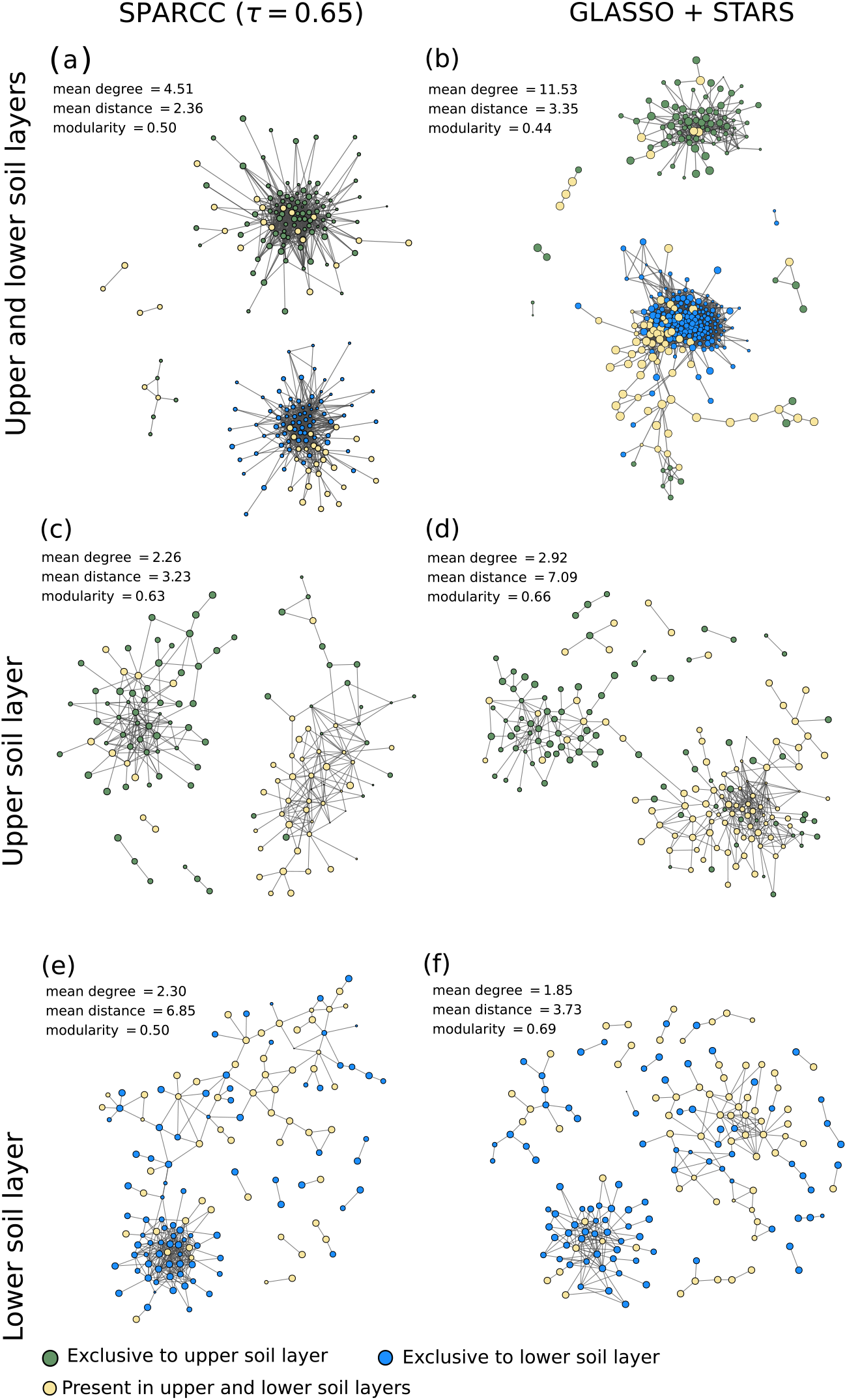
Networks of positive associations for microorganisms in a Beech forest soil at different soil depths constructed by different network inference methods. Networks constructed from upper and lower soil layers combined are shown in (a, b), and from each soil layer separately in (c-f). Networks were constructed by (left panel) SparCC or by (right panel) SPIEC-EASI-like approach (transformed with clr, and then using GLASSO combined with Stars). To improve visibility nodes with no edges are not shown.

**Figure 6:**
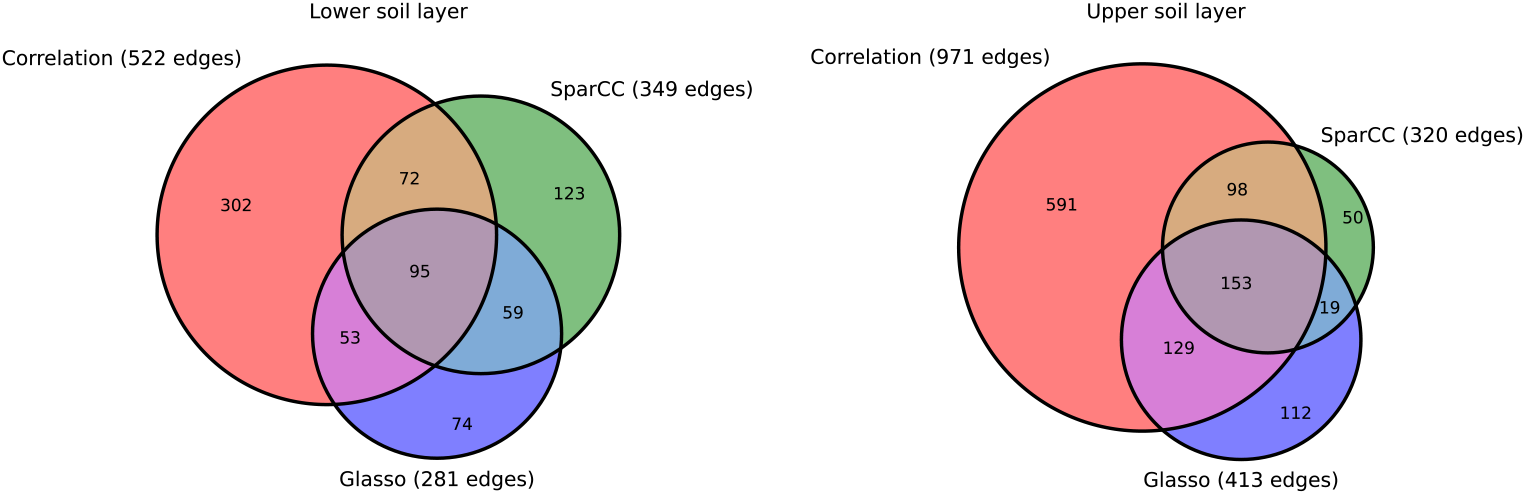
The edges detected by different network construction methods, using Spearman correlation (edge threshold 0.7), SPARCC (threshold 0.65) and GLASSO (STARS for model selection) with and the overlap among them.

### 3.3. Summary of test dataset analysis

We used microbial co-occurrence data from upper and lower soil layers of a European Beech forest to test whether the choices for a specific network inference method (i.e. correlation-based methods with different edge thresholds, (Fig. 4), a method using log-ratio transformations to address a possible compositionality bias, SPARCC, and a method considering indirect associations, GLASSO and Stars; (Fig. 5)), or the inclusion of an environmental factor (soil depth) would influence the structure of the constructed network, and the conclusions that can be drawn from it. The datasets of upper and lower soil layers together consisted of 466 ASVs, from which 162 were exclusively occurring only in the upper soil samples, and 183 only in the lower soil sample, and 121 were shared between the two (Fig. S5). We used this taxon-specific information to investigate if and to what extent the resulting network structure was influenced by taxon preference for upper or lower soil, two habitats which differed in many environmental conditions, like temperature, moisture, or substrate input. To visualize this, we categorized the nodes into those which were exclusively occur in one of the soil layers after filtering ASVs with low read numbers (Fig. S5), which could be interpreted as ‘specialists’ and those which were occurring in both soil layers, which could be interpreted as ‘generalists’. Irrespective of the method used, networks which were constructed including the environmental gradient, i.e. based on the combined dataset from both soil layers, always exhibited two highly connected sets of nodes (clusters), one containing all taxa dominant in the upper and the other those from the lower soil layer. This demonstrates that taxa which co-occurrence due to similar environmental preferences strongly dominate the network structure, potentially obscuring other ecological relationships.

Ruling out this environmental factor, i.e. constructing networks for each layer separately, revealed an interesting pattern. Here again we see in all methods that two clusters (modules) were forming, of which one was dominated by the specialists and the other one dominated by the generalists. This is an interesting finding, and the next step would be to think about its interpretation, or to formulate a testable hypothesis for future research based on this result. Both, however, go beyond the scope of this paper. Here, we just would like to point out that this pattern was conserved, in principle, in all the methods (Fig. 4, 5). The only observable limitation is that in the most sparsely connected network, i.e. the correlation network with a high edge threshold in the lower soil layer (Fig. 4f), this finding may not be visible anymore owing to the small number of connections. While the coarse pattern seems to be robust and independent of the method, however, we also observed that the majority of links were different between the different methods (Fig. 6). For a more detailed analysis, where pairwise associations between individual taxa are of interest, one should thus consider that methodological choices could influence conclusions at that level. For a display of networks with nodes coloured according to their phylogeny see (Fig. S7 and S8). Interestingly, however, both correlation (τ = 0.7) and SPIEC-EASI like methods identify the same central taxa that intermediated the connection between the modules of the upper soil layer (ASV_32a_pom, a *Proteobacteria Rhizobiales*). From this small test we conclude that strong patterns in the dataset will be captured by most methods, while the difference in the detailed connection pattern may be quite different, and conclusions on individual connections should thus be done with care.

## 4. Network Analysis

Network analyses can be applied to different types of data, f.e. data of soil samples collected in the field, root samples collected in different experimental settings, or even datasets generated by simulations (Ofaim et al. (2017); Hewavitharana et al. (2019); Berry and Widder (2014)). These datasets can create a variety of different networks such as resource competition networks (species and resources as nodes), food-webs (antagonistic interactions), mutualistic networks (positive interactions) or general co-occurrence networks. In this section we will illustrate the properties that can be central to formulate testable hypotheses for the biological system, see (Fig. 7).

**Figure 7:**
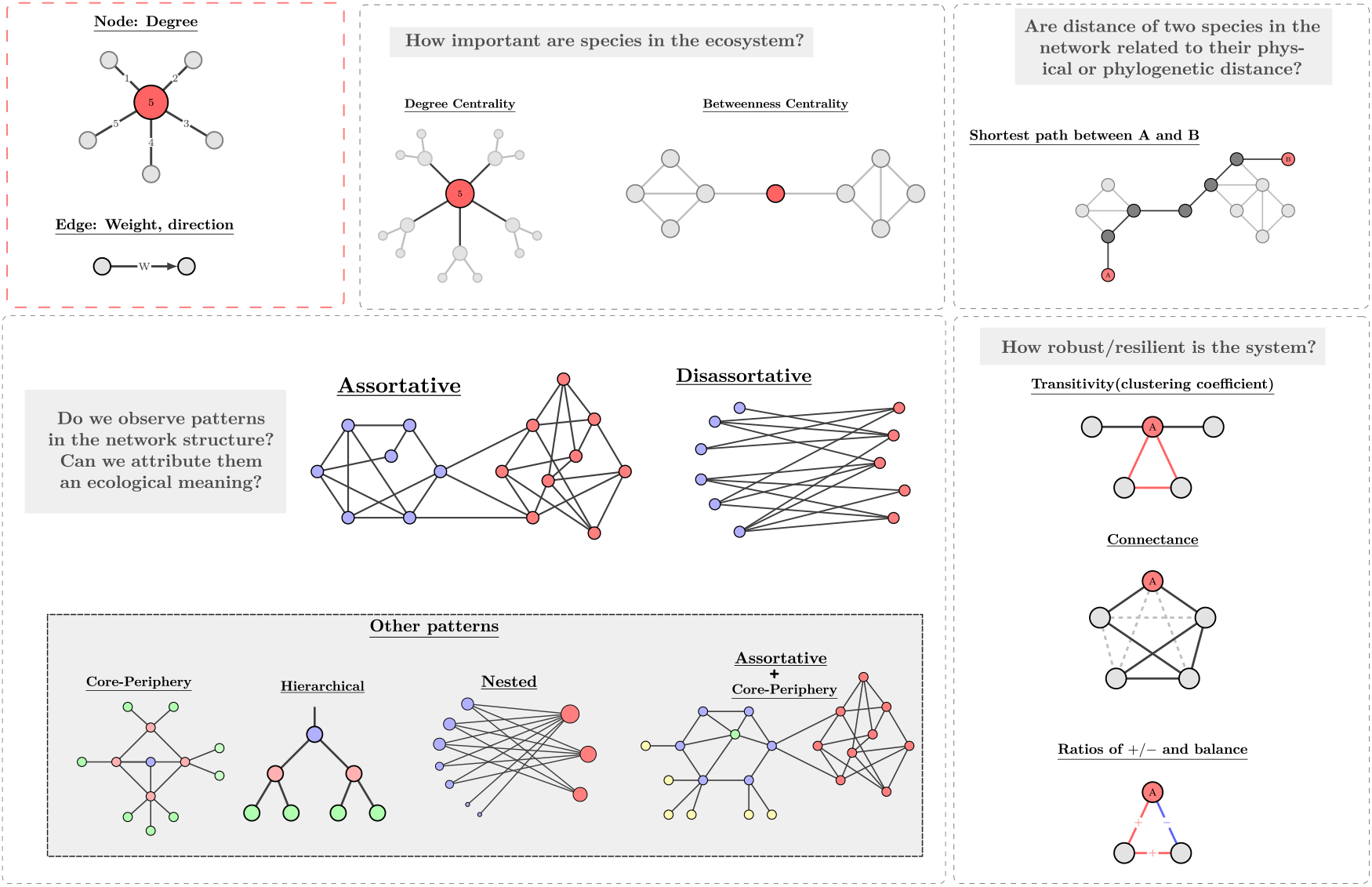
Some network properties that can be used to characterise microbial communities, with the respective examples of questions that can be addressed by their use.

### 4.1. Properties of the nodes

The number of significant associations and the positions of the node within the co-occurrence or interaction network are often used as benchmarks to assess key-stone species (Fig. 7). Keystone species are defined as taxa which play a pivotal role in the functioning of the ecosystem (Banerjee et al. (2018)), for example species which control fluxes of biomass (predators in food-webs). Identifying keystone species within a community is challenging (Power et al. (1996)). They can be detected empirically by removing individual taxa from the community and investigating the consequences for the functioning of the ecosystem. However, such experiments are difficult to carry out for soil microbial communities. As a keystone taxa is believed to have particularly many or special interactions with other taxa, it has been proposed that the number of edges or the position of a taxon within a network of interactions indicate its importance for the ecosystem (Banerjee et al. (2018)). While network analysis offers a novel way to look for potential keystone species by exploring node properties in co-occurrence networks, it has been emphasised that this can only serve as a method to suggest candidate keystone taxa. Whether they really occupy this role in the ecosystem needs to be validated empirically (Banerjee et al. (2018); Röttjers and Faust (2019); Banerjee et al. (2019)).

Using network properties such as node degree can thus be a valuable first step to formulate hypotheses to identify potential candidates for keystone species. The relevance of the node, however, is not only given by its degree, but also by its position within the network topology. This has been shown in model systems (gLV, generalized Lotka-Voltera), where the influence of the species is determined not only by its direct but through its indirect interactions (Berry and Widder (2014)). For this reason additional measures to identify the potential keystone species may be used. This can be done by determining the centrality (f.e. closeness, betweenness) of the node, see (Fig. 7) and (Tab. 2). Further improvement can be achieved by the knowledge of the topology and cluster structure of the network, see Sec. 4.3.

Despite difficulties, characterization of keystone species was successfully realized for some systems (predominantly gut microbiome). Using *Arabidopsis thaliana* as a model plant (Agler et al. (2016)) identified keystone taxa that establish a link between this plant and other microbes and have a central role in the community formation. For the soil microbiome, however, due to systems complexity, the concept of keystone species is not well established. It is speculated that microorganisms with a specific, but crucial function, such as nitrogen fixation or sulfate reduction (Pester et al. (2010)), can have strong influence without dominating the communities. In this context, taxa with strong enzymatic activity and decomposition ability were observed to also exhibit a high number of connections in co-occurrence networks (Zheng et al. (2021)). In agroecosystems microorganisms with a high number of associations in co-occurrence networks, f.e. genera such as *Actinobacteria* and *Proteobacteria* have been pointed out to have a great influence on the health of soils of different environments (Banerjee et al. (2018); Lupatini et al. (2014)).

### 4.2. Average path length of the network and degree distribution

The architecture of biological networks is often far from random. In general, interaction networks, such as food webs and mutualistic networks are known to have broad degree distributions, with few very connected nodes (hubs) and the majority of species that interact only with few others (Camacho et al. (2002); Montoya et al. (2006)). This degree distribution also leads to short path length between nodes, meaning that randomly chosen species have only few edges between them (Williams et al. (2002)). The questions are how and why such types of architecture develop for microbial communities, and their relation to stability or community assembly mechanisms. In the case of co-occurrence networks there is need to disentangle possible reasons behind the observed patterns, such as evolutionary history, dispersal dynamics or negative/positive biotic interactions (Morueta-Holme et al. (2016); Weiher et al. (2011); Goberna et al. (2019)). Relevant questions that still have to be approached are for example: how relative distances between taxa in the network are related to either phylogenetic similarity or even to physical distance if environmental gradient is present.

### 4.3. Cluster (network community) detection

Ecological networks can be composed of hundreds, or sometimes even thousands, of nodes. It is natural, therefore, to wish for identification of representative high-level subsets or groups to organise this complex information. However, we do not want any partition, but the one that can inform us of underlying functional, evolutionary or environmental units within these ecosystems. This is an easy procedure if we have metadata on such groups beforehand, f.e. division of microorganisms in groups predominant in certain environments ((Eldridge et al. (2020)). However, in most cases we lack this information and the underlying partition is precisely what we would like to learn about the system. The problem therefore is to determine node groups based on network topology, for instance by checking the density of connections among them (Newman (2010)). This is known as cluster detection or community detection. Note that this term is used for ‘network communities’, which consist of several similar nodes, not to be confused with microbial communities. This task consists of: (1) finding a natural division in the network, without being able to look at its structure; and more importantly (2) evaluating if these patterns are statistically/ecologically meaningful or if they can appear by mere chance.

Certainly the most popular algorithm developed for cluster detection (1) relies on modularity maximization, which formulates the task of community detection as an optimization problem. This is done by using a mathematical function known as modularity to assign a score to each division of the network (Newman (2004, 2006); Newman and Girvan (2004)). Although the method has attracted substantial popularity due to its simplicity, it has drawbacks which are often ignored in applications. While the method works well when a clear division exists, it is also known to find spurious partitions when there are none to be found, f.e. in completely random networks (Guimerà et al. (2004)). This conflicts with our objective (2) of selecting only statistically meaningful partitions. Moreover, a comparison with a null model cannot solve issue (2) since it can only rule out or accept the null model (random network structure), but cannot assess if the found partition is in fact statistically significant. This is because finding enough evidence to reject a null model is a much weaker requirement than having enough evidence to accept a particular network partition ^1^. Due to these problems several other methods have been proposed to extract meaningful communities from networks (Fortunato (2010); Fortunato and Hric (2016)). We highlight the inference methods based on stochastic block models (SBMs) (Karrer and Newman (2011); Abbe (2018); Peixoto (2019)). Their main advantages is their ability to distinguish signal from noise, taking care of (2) in a principled manner, and possess also the ability to detect not only patterns based on assortative structures (Newman (2004)), but others, such as core periphery, or self-similar hierarchical patterns (Fig. 7). The kinds of patterns identified by cluster detection have been proved useful for understanding ecological networks: Structural assortative as well as nested patterns have been linked to stability in interaction networks (Grilli et al. (2016); Thébault and Fontaine (2010)). In association networks clusters have mainly been used to identify and separate different niches occupied by microorganisms.

### 4.4. Network stability

Over the past decades an increasing number of studies have approached the question of ecosystem stability by analyzing the architecture of biological interaction networks (Landi et al. (2018); Montoya et al. (2006); Bascompte (2009); Okuyama and Holland (2008)). The stability is characterized by how systems respond to disturbances, and whether or not they can return back to their original function after being perturbed. To evaluate stability one needs to estimate either the recovery process (resilience), endurance (robustness) or the flexibility of the system (persistence) (Landi et al. (2018); Hodgson et al. (2015)).

A significant fraction of the studies on stability are theoretical and use numerically generated datasets (f.e. gLV, generalized Lotka-Voltera model). They address a long standing debate in ecology of the relationship between the ecosystem’s complexity (f.i. number of connections in an ecological network) and its stability. The controversy in the field started around the 1970s when theory showed a conflict between the two aspects (May (1972); MAY (1974); Sales-Pardo (2017)). Currently we know the network properties that foster stability, but only for some types of networks. For example, nestedness and connectivity seem to promote stability in mutualistic (Thébault and Fontaine (2010)) and resource competition networks (Wei et al. (2015)), while for food-webs a compartmentalized structure, low connectance seem to bring stability (Gross et al. (2009); Neutel et al. (2007)). At the same time, evaluation of the stability of an ecosystem from the structure of co-occurrence networks, where the role of links is less clear, is only at its infancy. In general what is speculated is the presence of mutualistic interactions may destabilize ecosystems, since they generate co-dependencies (Coyte et al. (2015)). Trophic interactions, on the other hand, can promote stability due to negative feedbacks (Coyte et al. (2015)). Finally, the level of clustering and the degree distribution in the network are other very important factors that affect the stability of ecological communities, (Grilli et al. (2016)).

Studies that compare the structure of co-occurrence networks from (experimental) ecosystems under natural and disturbed conditions help to reveal the potential link between network structure and system stability. Interesting examples of such are (Yuan et al. (2021); de Vries et al. (2018); Wagg et al. (2019)). Some of such studies already have shown that connectivity, density of the links and transitivity decrease in disturbed environments (Karimi et al. (2016, 2017)). It is, therefore, expected that these properties could be used to assess the response of microbial communities to changes in their environment. In addition, formulation of hypotheses on the resilience of a system can be an important way to validate network models inferred by the current methods. For example, tracking the presence of identifiable keystone species and how the network structure is affected by them can be of interest. Finally, we would like to mention the appeal of using co-occurrence networks as indicators of ecosystem’s quality and function as proposed by (Karimi et al. (2017)).

## 5. Challenges and way forward

### 5.1. Specific challenges in soil microbial ecology

Although there are several possible drivers behind the formation of microbial communities, species-species interactions have taken the spotlight of current research on taxa co-occurrence patterns. Despite its appeal, the detection of interactions from this type of data has proven to be extremely challenging. Moreover, it is still unclear whether our limitation in spotting species-species interactions can be attributed to an inefficiency of current inference methods, or to the weakness of interaction signal in abundance (co-occurrence) data (Blanchet et al. (2020)).

Clearly, besides the potential signal of species interactions, association networks contain other valuable information about relationships within an ecosystem and can help us to understand ecological processes behind the high microbial diversity characteristic to soil. However it is important to emphasize that any network model and its analysis is only as good as our interpretation of it. Therefore, it is desirable to include an interpretation to all associations represented as edges to the most possible extent. This is particularly hard for soils, since they are characterized by their immense biological diversity and heterogeneous physical structure. Here we summarize the important soil characteristics that may impact network construction, interpretation and analysis, to then next discuss how we can use experimental design and statistical methods to improve our ability to reconstruct ecological relationships.

#### 5.1.1. Soil microbial datasets, what to include as nodes?

Soil harbours one of the highest biodiversity of all microbial environments on Earth as was observed in the Earth Microbiome Project (EMP; https://earthmicrobiome.org/) (Thompson et al. (2017); Walters and Martiny (2020)). The characterization of this diversity is not easy, and depends on the number of samples used and the design of the sequencing experiment (Hermans et al. (2019)). Increasing the sampling effort by including more replicates results in discovery of larger numbers of rare taxa. Although it is considered that these taxa significantly contribute to ecosystem processes (Hermans et al. (2019); Jousset et al. (2017)), it is still unclear how they can be included in the network construction, as taxa with low prevalence lead to spurious associations, and are thus often completely discarded in the data preparation step (see Sec. 3.1).

Fungal-bacterial associations are another important component of the soil ecosystem (Peay et al. (2016)). The understanding of fungal diversity across biomes remains more limited due to challenges in sequencing the ITS region of fungal DNA. Integrating the community sequence data from fungi and bacteria to construct combined networks will certainly expand our ability to explain the observed community patterns in the soil environment. However, the construction of such cross kingdom networks faces new challenges, since the analysis would have to properly combine two compositional datasets. Despite this, cross-kingdom networks have been constructed by applying correlation-based approaches on merged 16S rRNA gene and ITS relative abundance datasets (Delgado-Baquerizo et al. (2020); Jiao et al. (2020)). We would however argue against such an approach, as combining correlations within and across compositional datasets in the same network may lead to a spurious connection pattern. Possible alternatives propose the use of centered log ratios individually on each dataset before merging them together (Tipton et al. (2018)) or transform the relative abundance data into a presence/absence dataset (which implies a loss of information). In addition, one can, depending on the research question, consider constructing bipartite networks, which only contain edges between the kingdoms (Montesinos-Navarro et al. (2012); Toju et al. (2015), or multilayer networks, which contain different types of edges, (Pilosof et al. (2017)).

### 5.2. Soil microbial networks: what type of associations should we expect?

Soil provides a unique environment for microbes, because it is spatially structured and chemically and physically heterogeneous across several scales. Within the complex soil matrix, numerous edaphic properties impact the structure of the microbial communities therein, including pH, salinity, temperature and moisture (Zheng et al. (2019); Frindte et al. (2019)). Additionally, temporal dynamics such as weather patterns, root exudation and other seasonal inputs of organic material may influence the structure and activity of a microbial community (Kuzyakov and Blagodatskaya (2015); Chernov and Zhelezova (2020)). As a result of soil heterogeneity and temporal fluctuations, it is critical to consider that sequence-based approaches reveal only a snapshot of the microbial community present at a given time. Together, these properties of soil as a habitat for microbes influence the possible scope for interpretation of microbial association networks.

Heterogeneity of soil samples is a major source of concern in the development of reliable inference methods to uncover ecological relationships. We see the heterogeneity impacting the sampling and inference in two different ways. First, it can lead to hidden physical or chemical differences between apparently similar samples (Armitage and Jones (2019)). Since environmental drivers lead to strong differences in the community structure, such drivers may dominate the detected association signal. In this case, additional environmental measurements can help to interpret the observed patterns. The second concerns the large volume of each sample used in the analysis. Nucleic acids extracted for sequencing studies are derived from amounts of soil ranging from 250 mg to 2 g of material. Very often, they are aliquots from an even larger soil volume, for example several soil cores that have been mixed and homogenized to be representative for a certain field site or plot. Soil microbes, however, interact with each other at the scale of tens of *µm* (Raynaud and Nunan (2014)). It is obvious that co-occurrence of microbial taxa across soil samples taken at a scale which is by orders of magnitude larger than the scale at which they interact, does not necessarily mean that they interact with each other. In line with this, community assembly theory suggests that the larger the sampling units scale the more environmental filtering effects dominate species co-occurrences (Kraft et al. (2015)). Although the mix of ecological factors operating at larger sampling scales obscures causal distinctions, studies have shown that networks generated with this type of data hold predictive value for central ecosystem functioning (Shi et al. (2016); Wagg et al. (2019)). We should, however, lower our expectations to capture a clear signal of microbial interactions, since even small soil samples may contain numerous metacommunities – small consortia linked by dispersion and diffusion (Armitage and Jones (2019)). Two microbial taxa can be separated by unconquerable distances even in a small piece of soil given its microscale physical structure and the spatial heterogeneity of microbial niches. In view of this complexity it is controversial if in such samples biological interactions can leave any signal at all. For example, models suggest that negative interactions, such as competition, can be completely hidden at scales much larger than the scales of the interactions themselves (Araújo and Rozenfeld (2014)). In summary, the associations described in co-occurrence networks from standard soil samples have to be carefully interpreted. Such networks are expected to be dominated by associations driven by the environment, however the obtained network structure may be a result of several unknown factors. In relation to interactions among organisms, especially the signal of competitive interactions will be obscured. In the next section we give suggestions on how to disentangle the different factors behind a given co-occurrence dataset and suggest alternative or complementary experiments.

### 5.3. Reconstructing networks of ecological relationships

#### 5.3.1. Network interpretation is influenced by experimental design

In general, edges in networks based on the characterization of variability of microbial communities in multiple soil samples can inform about species co-occurrence patterns across these samples. If the objective is to infer microbial interactions, we advise to work with datasets extracted from specific micro-environments where interactions are prone to occur. Such environments can be, for example, mycorrhizal fungi within plant species root tips (Montesinos-Navarro et al. (2012)), soil aggregates (Szoboszlay and Tebbe (2021); Wilpiszeski et al. (2019)) or synthetic systems (Cordero and Datta (2016)). In the case of soil aggregates, it is possible that niche partitioning also leads to spatial segregation with their pore structure. However, even in this case, aggregates are considered to provide enough physical proximity and a traversable matrix necessary for competitive and faciliatory exchange. This is critical as aggregates contain microbes’ primary source of nutrients and their occupancy enables the establishment of stable source populations. Alternatively, to improve interpretation one can amplify and sequence specific functional genes in place of taxonomic marker genes, to target microbial groups with known traits, f.e. as was done in (Jones and Hallin (2019)) to find relationships between nitrifiers in soil. Certainly, in networks based on functional information, such as metabolic profiles associated with distinct groups of microorganisms that transform metabolites, edges more likely represent potential interactions among taxa (Ofaim et al. (2017); Hewavitharana et al. (2019)).

Another important factor that influences our interpretation of edges is the distribution of the heterogeneity across samples. Microbial datasets collected along large environmental gradients can reveal differences in environmental niches occupied by microorganisms. Indirect associations through environmental factors will also be predominant if more than one treatment is considered within the same network. For example, datasets encompassing different seasons, depths, temperatures and biomes will be prone to generate co-occurrence patterns derived from the different levels of the treatment, but will also blur the patterns of co-occurrence among taxa within the levels of the treatment, as is also demonstrated in our example of microbial networks across different soil depths ((Fig. 4, 5), Sec. 3.3). Therefore, the inclusion (or not) of treatments in network construction will impact which ecological processes can be assessed. The focus of the hypothesis to be tested will define whether networks should be constructed from combined data of different treatments, or whether separate networks should be built for each homogeneous environment.

Several strategies were proposed to understand which one of the established edges are in fact due to the indirect influence of environmental factors (Faust (2021)). A strong imprint of environmental factors or treatments often leads to clusters or modules being assorted by these factors. This can be tested for by categorizing taxa according to the environment or treatment where they predominantly occur and check if the network structure is shaped by these categories (Jiao et al. (2020), (Fig. 4, 5)). If there is an obvious difference between treatments or environmentally different sites, depending on the research question, these could be compared and studied separately. In that way, the influence of the environment on the network structure would be reduced, allowing other potential ecological factors, such as interactions, to become more visible in the network structure. Alternatively environmental factors can be directly integrated into the network construction, for example by representing them as nodes in the network, enabling one to examine their influence on the other nodes (taxa) (Faust (2021)). In the same way, data on the distance between samples can also be used to check if the links result indirectly due spatial dispersion patterns (Goberna et al. (2019); D’Amen et al. (2018)).

#### 5.3.2 Network interpretation can be improved by statistical methods

In summary edges in a network construction can appear due to: (1) species-species interactions, (2) association mediated by the environmental factors (niche overlap), (3) spatial variability due to dispersion dynamics, (4) due to influence of a third interfering species present in the network (common to methods based on correlations), (6) spurious associations due to inappropriate data handling (errors in dealing with sparsity and compositionality) and (7) noisy measurement. It is clear that the possible interpretations of an ecological network rely not only on the data but also on the statistical methods used for its construction, especially with respect to items from (4-7) listed above. As we have described in Sec. 3 these methods can significantly differ in their assumptions and strategies, and therefore a single dataset can produce networks with very distinct sets of edges, see (Fig. 6). Part of the difficulty current approaches meet is addressing the nature of amplicon-sequence data itself (item 6), we advise the use of qPCR to reliably infer the abundances of taxa from sequence read numbers (Alteio et al. (2021)). We emphasize again here that correlations are unreliable to establish interpretable edges, since they do not offer a clear criteria for distinguishing signal from noise and to distinguish direct from indirect associations, see Sec. 3. This can be improved with model-based inference approaches and incorporation of additional information (environmental factors, distance between samples) into the network construction can strongly facilitate edge interpretation (items 1-3). At the moment, the most promising direction is biologically relevant generative models, such as those based on generalized-Lotka-Voltera equations (gLV) Bucci et al. (2016); Gibson and Gerber (2018). Although these methods can be computationally demanding and are still not sufficiently efficient for large data sets, this is a very rapidly advancing area where alternatives constantly appear.

#### 5.3.3. Questions that can be addressed in different networks

The nature of the edges in a network affects the type of questions that can be addressed with network analysis. Ecological questions can be assessed with network analyses when particular network properties match the expected ecological patterns. Thus, when it is expected that groups of taxa tend to interact more among them than with other taxa, network properties such as network community structure (modularity) can be useful to test that specific hypothesis. In some cases, a given network property can be informative, independent of whether the edges represent interactions or only have a more broad meaning such as co-occurrence (including niche overlap). In that sense, it might be interesting to test whether there are groups of taxa that tend to interact or that tend to co-occur. However, other network properties can only meaningfully be evaluated only when we know that edges describe interactions among taxa. For instance, that might be the case of the average path length among nodes, which can reflect the contribution of complex indirect (i.e. high average paths length) vs. direct interactions among taxa (i.e. short paths), but has a less useful interpretation in terms of co-occurrence. As explained in the previous subsections the experimental design from which the data is obtained and the statistical methods used will constrain the meaning of the edges in a network, which in turn will condition the network properties that would be useful to assess specific ecological questions (Table 2).

**Table 1.**
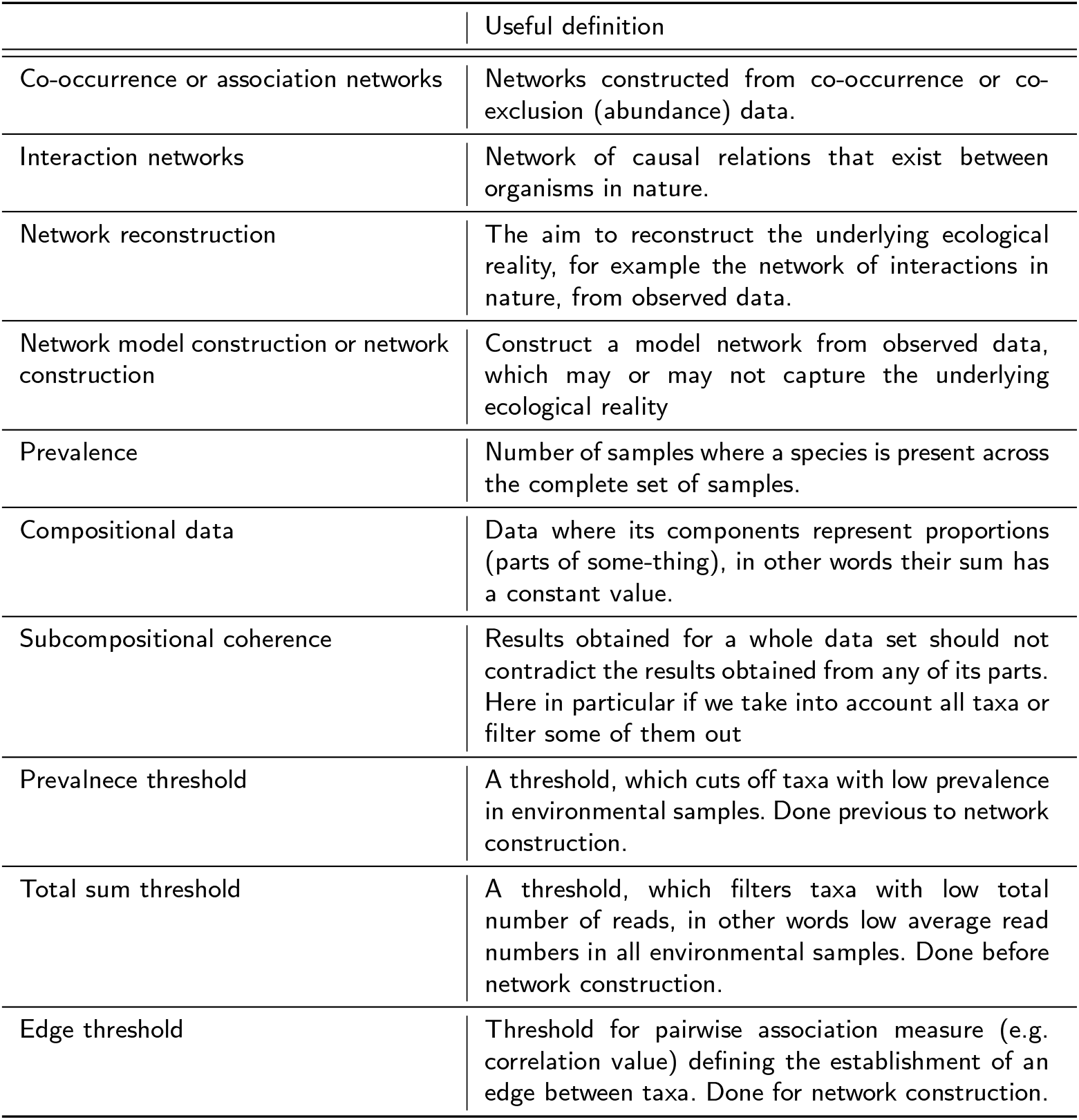
Definitions used in this Perspective Paper.

|  | Useful definition |
| --- | --- |
| Co-occurrence or association networks | Networks constructed from co-occurrence or co-exclusion (abundance) data. |
| Interaction networks | Network of causal relations that exist between organisms in nature. |
| Network reconstruction | The aim to reconstruct the underlying ecological reality, for example the network of interactions in nature, from observed data. |
| Network model construction or network construction | Construct a model network from observed data, which may or may not capture the underlying ecological reality |
| Prevalence | Number of samples where a species is present across the complete set of samples. |
| Compositional data | Data where its components represent proportions (parts of some-thing), in other words their sum has a constant value. |
| Subcompositional coherence | Results obtained for a whole data set should not contradict the results obtained from any of its parts. Here in particular if we take into account all taxa or filter some of them out |
| Prevalence threshold | A threshold, which cuts off taxa with low prevalence in environmental samples. Done previous to network construction. |
| Total sum threshold | A threshold, which filters taxa with low total number of reads, in other words low average read numbers in all environmental samples. Done before network construction. |
| Edge threshold | Threshold for pairwise association measure (e.g. correlation value) defining the establishment of an edge between taxa. Done for network construction. |

**Table 2.**
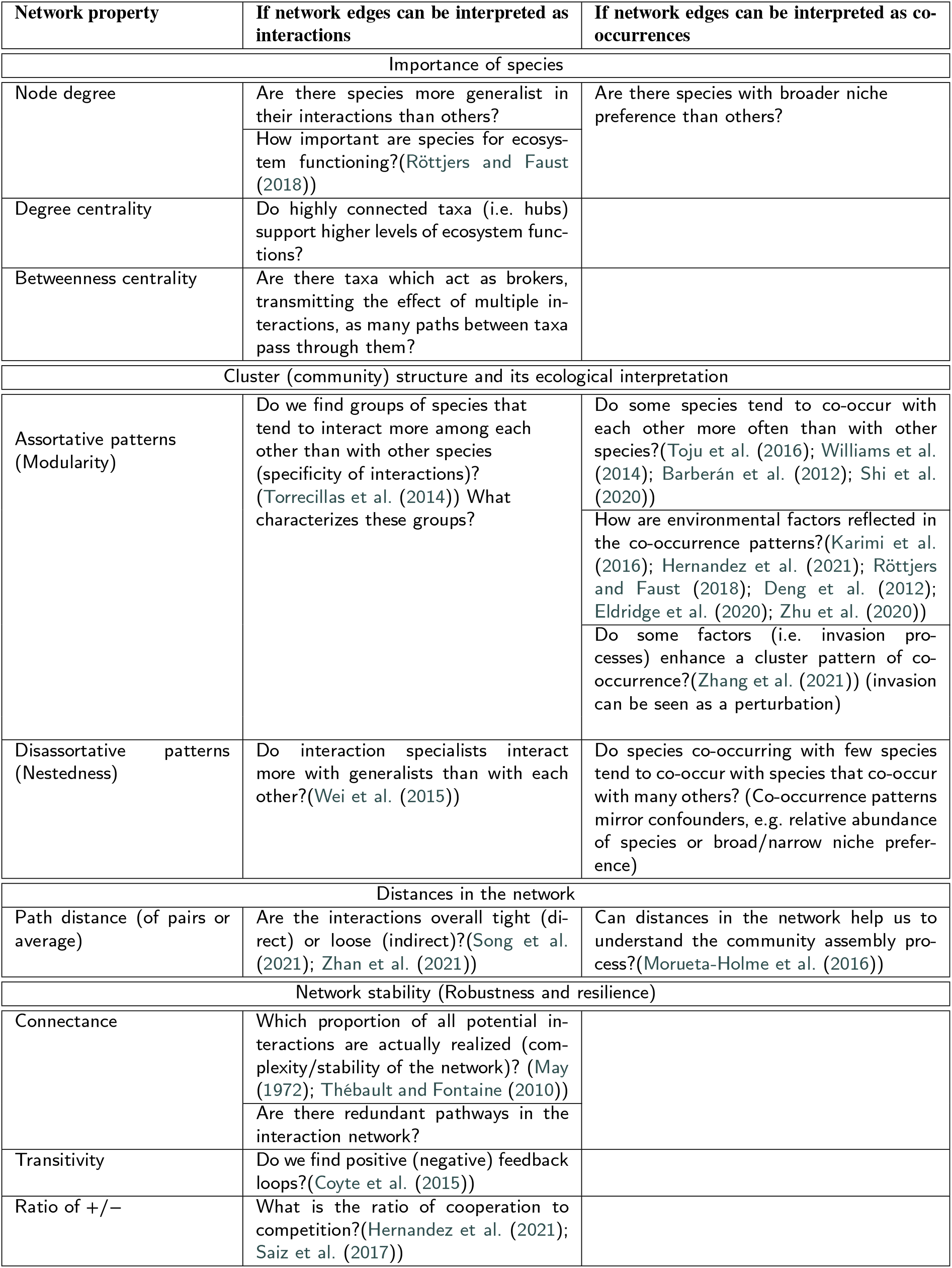
Examples of ecological questions that can be assessed with particular network properties depending of the meaning of the links in the network.

#### 5.3.4. Descriptive, exploratory or hypothesis-based approaches?

Recently we have observed an increase in the number of articles which use microbial networks as a form of data visualization for co-occurrence patterns in soil. These studies define the edges of networks as ‘co-occurrences of microorganisms’ (or even incorrectly as microbial interactions), and have a mainly descriptive character. Without presenting hypotheses of biological mechanisms behind the observed statistical patterns in co-occurrence data, such studies are of little interest to soil microbial researchers. In the previous section we presented several research questions which can be addressed with network analysis. Addressing these questions using the constructed network can be a simple form to extend and improve such descriptive studies. Networks should work as a tool to help us understand the processes that structure microbial communities, and therefore the interpretation of observed patterns and creation of hypotheses from these patterns are not optional and should be included.

Finally, we shortly discuss possible ecological hypotheses that might be tested or used in microbial network analysis. As described in Sec. 4.1, identification of potential keystone species or functional components of the network can help us to forecast structural shifts in the network in response to environmental perturbations, see (Fig. 8c). Another possible research direction is to compare the network architecture shaped by particular conditions, f. e. the available substrate, see (Fig. 8c). In this context, networks can appoint taxa with potential similar functionality, since those are expected to compete and be negatively associated across communities (Brown and Wilson (1956)). In the same way we can formulate hypotheses on functionally complementary species, expected to be present in the degradation of complex substrates (Lindemann (2020)). This can be done using networks constructed from datasets extracted from different environmental conditions, expecting a change in predominance of facilitation to competition with increasing nutrient availability (Hoek et al. (2016)). These nutrient-based community responses can be combined with a stress response (stress imposed by an environmental factor). What is expected then, is that high stress would reduce investment into competitional traits, while facilitation would enable stress mediation, equivalent to what we know from plant ecology (Bertness and Callaway (1994); Hammarlund and Harcombe (2019); Piccardi et al. (2019)).

**Figure 8:**
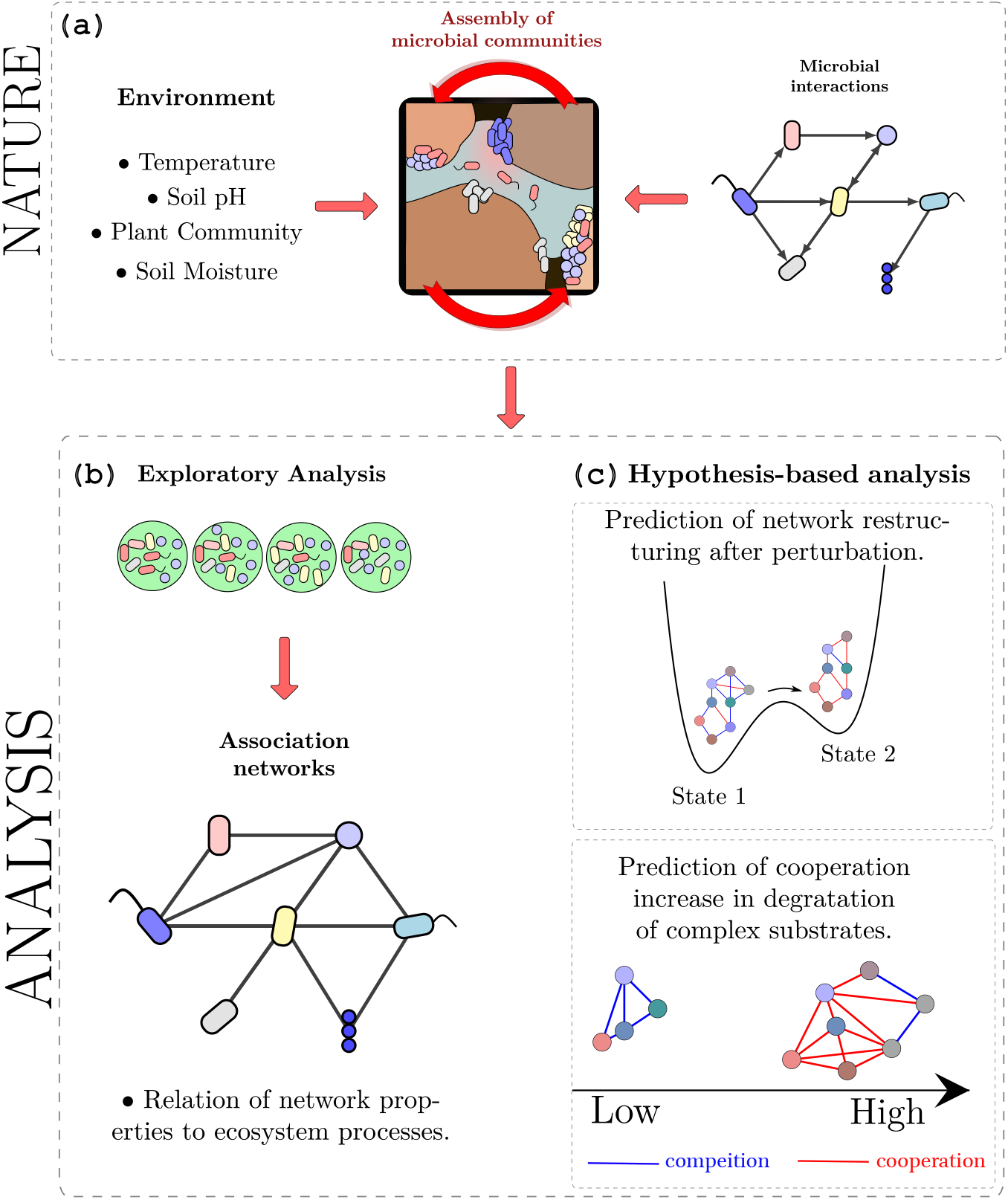
Use of network analysis in exploratory vs hypothesis driven research. (a) Taking into account that community composition is shaped by underlying ecological mechanisms driven by the physical environment or by interactions among taxa. (b) In general, the measured co-occurrence patterns are used to construct association networks, which are different from the interaction networks. (c) Hypothesis-driven research is key to design comparative experiments aiming to isolate and contrast the community assembly mechanism of interest. In particular, established ecological theories can complement hypothesis generation.

## 6. Conclusion

As our understanding of soil biological diversity has increased with the use of high-throughput sequencing techniques, the new challenge for the field of soil microbial ecology has become to understand the complex web of relationships among microorganisms. This remarkable shift in the field is reflected in a rising number of papers using network analysis. The central feature of network models is that they naturally bring microbial associations to the center of our attention. The approach forces us to see the taxa not as isolated units, but to build an integrated picture of microbial communities and also of their surrounding environment. Although these network models open a whole new dimension of possible research, at the moment we have to proceed with caution. The reason is that we are still lacking reliable inference methods and consistent experimental protocols that can successfully reconstruct the hidden network of ecological relationships from environmental samples. Therefore we can only grasp part of the potential of network analysis with a careful interpretation of results obtained from co-occurrence analysis. Finally, we emphasize that the potential of networks goes far beyond being a visualization tool and the information gained from networks constructed from soil microbial community data can – despite limitations – deliver valuable insight into ecosystem organization when applied in hypothesis-driven research. While the traditional question addressed by DNA sequencing analyses of soil microbial communities is “who is there”, the novel question which can be asked by network analysis, when successfully applied, is ‘who co-occurs with whom, and why?”.

## Acknowledgements

KS, LA, ES and CK have received funding from the European Research Council (ERC) under the European Union’s Horizon 2020 research and innovation programme (grant agreement No 819446).

## Footnotes

1 For example, a network partition where only one of hundreds of detected communities is statistically significant will be enough to reject a null model, but it would be mostly statistical noise.

## Notes

### Competing Interest Statement

The authors have declared no competing interest.

