## Supplemental Material for "From Diversity to Complexity: Microbial Networks in Soils"

### I. ILLUSTRATION OF THE INCREASE OF THE USE OF NETWORK ANALYSIS IN SOIL RESEARCH

In the last two decades we have observed a significant increase of the use of network analysis in the investigation of microbial systems. We illustrate this increase by showing the number of articles published in the field from 2000 to 2020, see Fig. S1. The figure shows the result of a keyword search in Web of Science, by selecting the research articles with expressions: “microbial network analysis” or “microbial co-occurrence analysis”, and filtering the ones applied to soil with expression “soil”. We can observe an exponential growth of such articles in the past decades, and the rate of increase is faster in soil research (see inset of Fig. S1). Having only less than 10% from the total until 2010, they have reached almost 35% of all microbial network papers by 2020.

### II. SAMPLING SITE AND PROCEDURE

Soil was sampled in a mature beech forest (48°07′18.0”N 16°02′45.8”E, 540 m altitude) after removal of litter, on the April 12, 2021. The sampling site is located at the research site ”Klausen-Leopoldsdorf” in the Vienna Woods, Lower Austria, Austria, which is run by the Austrian Federal Office and Research Centre for Forests. The soil type is classified as a dystric cambisol over sandstone with a loam–loamy clay texture [3]. Soil cores (8 cm diameter, 5 cm height) were taken at two soil depths ($0-5$ and $15-20$ cm) one on top of the other, at two locations with approximately one meter distance to each other in five plots, respectively. This made up 10 soil cores for each soil depth. The cores were transported to the laboratory and kept at 4°C overnight. Subsequently, we took two subsamples (core: 3 cm diameter, 5 cm height) from each core except for one core of the lower soil horizon, which contained too many stones to core it. We ended up with 20 sub-cores of the upper soil horizon and 18 for the lower soil horizon. After we sieved the cores (2 mm) and removed roots and

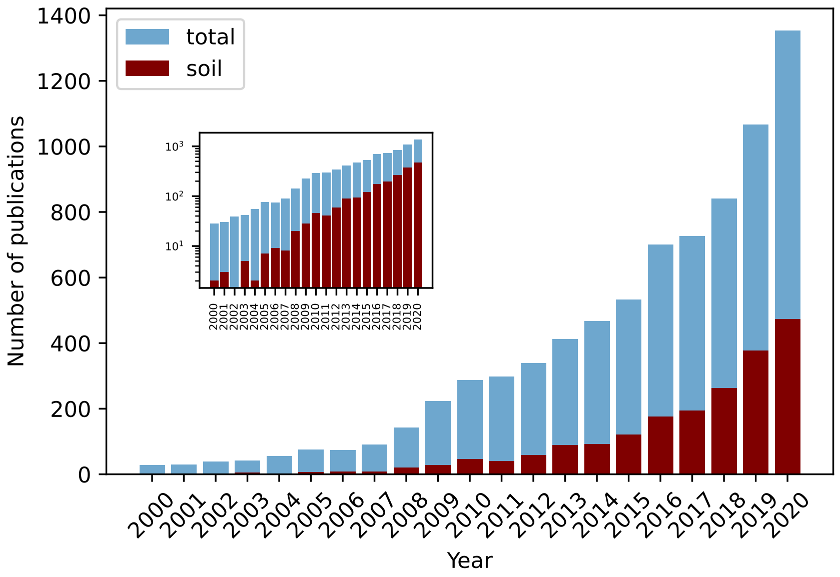


FIG. S1: Number of publications from a Citation Report from Web of Science using for the total the expression (((ALL=(microbial network analysis)) OR ALL=(microbial co-occurrence analysis))) and for soil the expression ((((ALL=(microbial network analysis)) OR ALL=(microbial co-occurrence analysis)) )AND ALL=(soil)). Inset: the plot in log-lin scale, to show the exponential growth.

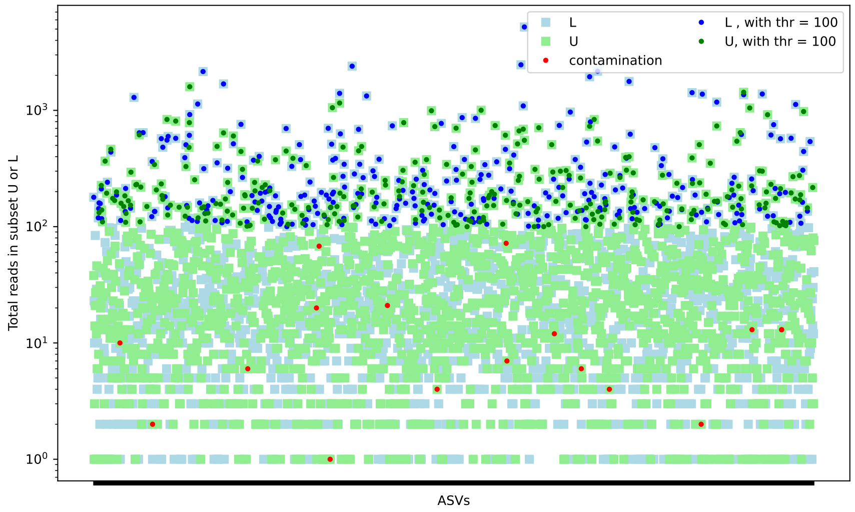


FIG. S2: The dataset has 38 samples, that can be divided in two subsets with 20 samples from the upper soil (U — light green) and 18 samples for the lower soil (L – light blue). The figure shows the total number of reads for each ASV (1899 in total) for each subset. Red dots represent the contaminants detected with Decontam [2]. For the network analysis we have selected only the ASVs with more than 100 total reads in each dataset. This criteria leaves us with 283 ASVs for the analysis of the upper soil (green), and 304 ASVs in the lower soil (in blue).

stones, we weighed in 400 mg of each of the 38 samples in Lysis Matrix E tubes (MP Biomedicals) and stored them at $-80°$ until DNA extraction.

### III. DNA EXTRACTION AND SEQUENCING

Collected soils ( 400 mg of soil) underwent bead-beating using Lysis Matrix E tubes (MP Biomedicals) on a PowerLyser (MP Biomedicals) for 30 seconds. Following bead-beading, DNA was extracted from the soil samples at the Joint Microbiome Facility (JMF) [5] using the MPBio Soil DNA extraction kit (MP Biomedicals). Extracted DNA was assessed for quantity and quality prior to PCR amplification using the 515F/806R primers for the 16S SSU rRNA marker gene. Amplified material was sequenced on the Illumina MiSeq platform (Illumina).

### IV. DATA PREPARATION

#### A. Sequence Analysis

Sequenced reads were assessed for quality and ASVs were inferred using DADA2 software by the JMF [1, 5]. Taxonomy of 16S rRNA gene sequences was classified using SINA (v.1.6.1, https://www.ncbi.nlm.nih.gov/pubmed/22556368) against the SILVA NR99 database (release 138.1, https://www.ncbi.nlm.nih.gov/pubmed/23193283). The sequences were classified into 1899 ASVs. Data analysis was conducted in R using the phyloseq package ([4]). Contaminating sequences were detected and removed using Decontam software [2], see red dots in Fig. S2.

The obtained number of reads for the samples originating from lower soil layer were similar to the ones from upper soil layer. The sparsity of the dataset is shown by the mean-variance relationship of the sequence reads in Fig. S3. For the network construction we do not use taxa with low read numbers (these taxa also correspond to taxa with low prevalence), see Fig. S2 and S3. This criteria leaves 283 ASVs in the upper and 304 ASVs lower layer. After trimming we leave only the ASVs with more than 100 total reads in each dataset (this also corresponds to an average of 5 reads per sample). The taxonomic composition at the Phylum level of datasets are shown in Fig. S4, the main differences among the subsets are the higher fraction of 16S rRNA attributed to *Actinobacteria* and *Bacteroidetes* in the upper soil layer, and higher fraction attributed to *Chloroflexi* in the lower soil layer. We show the overlap between the detected taxa in both layers in Fig S5, the two datasets share 121 ASVs.

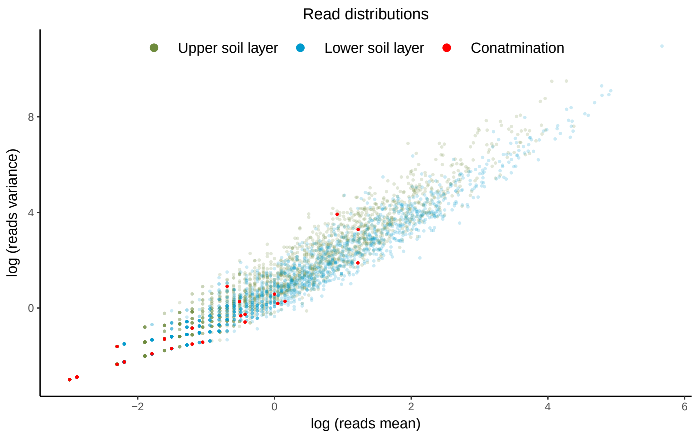


FIG. S3: Read distributions - means related to variance - for upper soil layer (in green) and lower soil layer (in blue). Contaminating sequences from either dataset are highlighted in red.

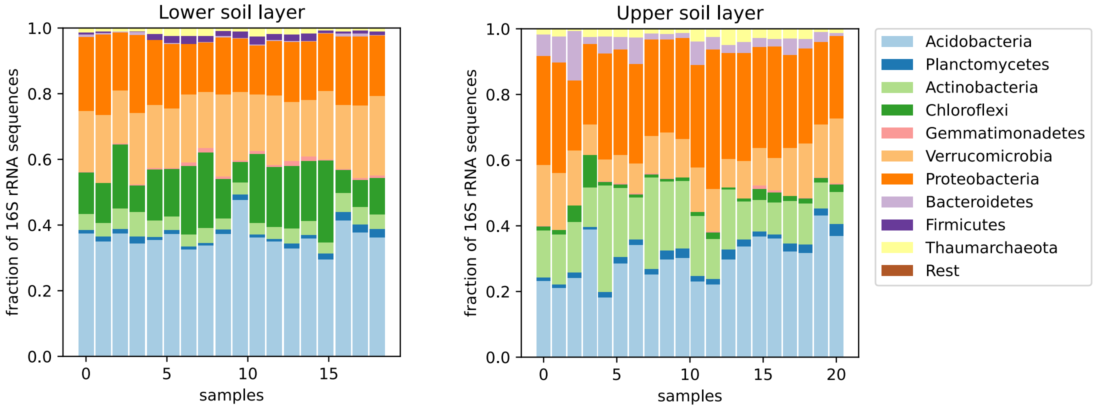


FIG. S4: Taxonomic composition of the communities in the two subset of samples after filtering (removal of ASVs with less than 100 sequenced reads).

#### B. Influence of zeros

As we write in the main text and show in Fig. S3, the high-throughput sequencing datasets have a large number of zeros. These zeros can have different origins and therefore may distort the network construction. In other words to infer associations on the basis of these zeros can be misleading. At the same time the presence of zeros results in strong pairwise correlations for rare taxa as shown in Fig. S6 (a, b). Fig. S6 (a, b) represent the edges detected for the non-filtered dataset, after an edge threshold of 0.7 for pairs of taxa with given prevalence in the dataset. Notice the grid of 20x20 samples (circles) for the upper layer, 18x18 for the lower layer. The figure shows that a higher number of edges is established between taxa with low prevalence. Filtering out the taxa with low prevalence, or taxa with low total read numbers strongly changes this pattern as shown in Fig. S6 (c, d). In general we see a lower number of edges established, and most of them are for prevalent taxa.

#### C. Constructed networks

The steps for the network construction are described in the main text. Here we reproduce the networks showing the phylogenetic classifications of nodes in Fig. S7, S8.

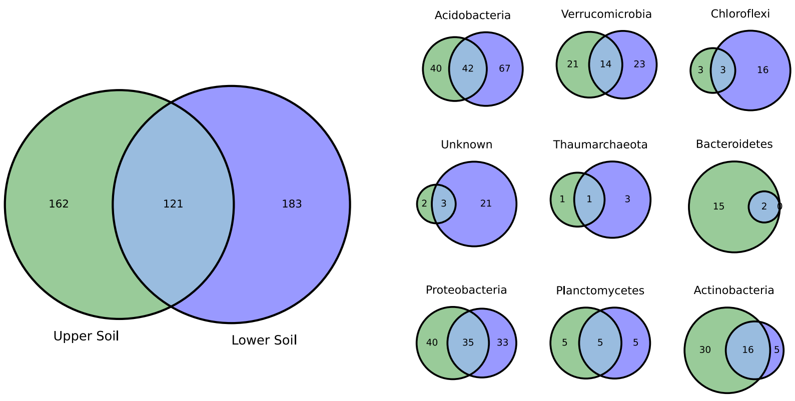


FIG. S5: After filtering the ASVs with low read numbers we are left with 283 ASVs for the upper soil layer and 304 ASVs for the lower soil layer. The two datasets after the filtering share 121 ASVs.

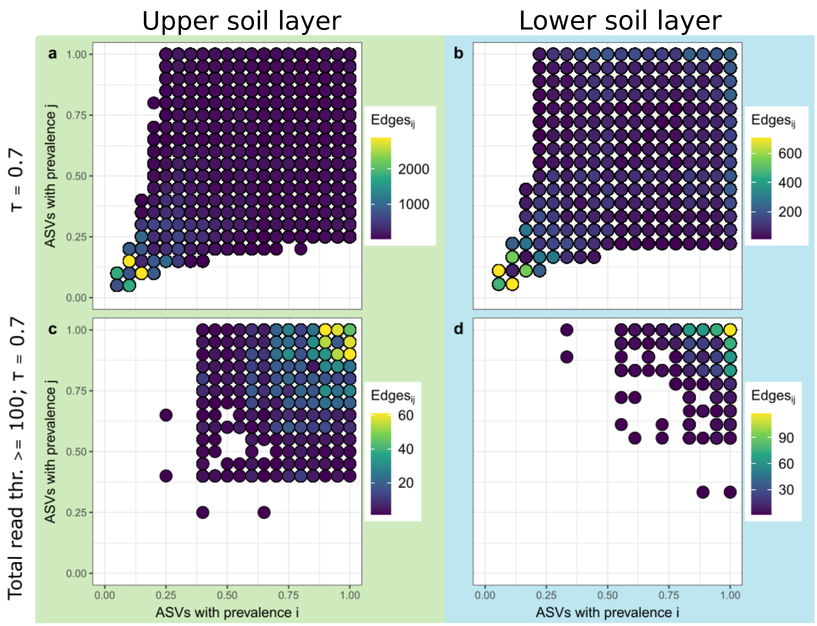


FIG. S6: Edge counts between pairwise associated ASVs plotted by either partners prevalence (i and j). Plots (a) and (b) represent the full datasets. Plots (c) and (d) represent filtered datasets, where ASVs with total read counts below 100 across all samples were eliminated. In both cases associations were established from Spearman correlations for relative abundances and coefficients were trimmed by a threshold of $\tau = 0.7$. Plots a) and c) show upper soil layer associations and b) and c) show lower soil layer associations.

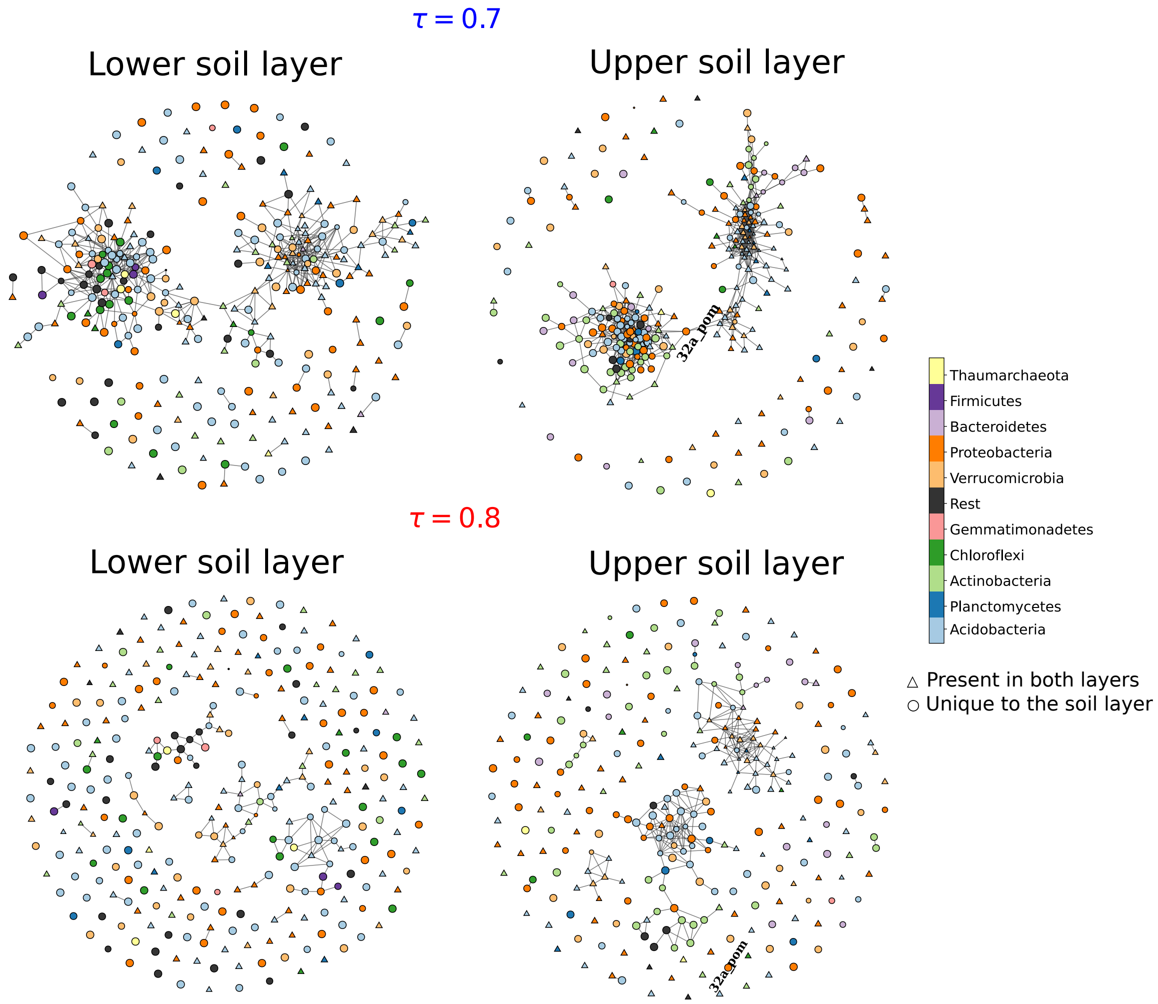


FIG. S7: Association networks constructed from Spearman's rank correlation coefficient, for $\tau = 0.7$ (upper panel) and 0.8 (lower panel) with phylogenetic classifications of nodes (complements Fig.3 of the main text).

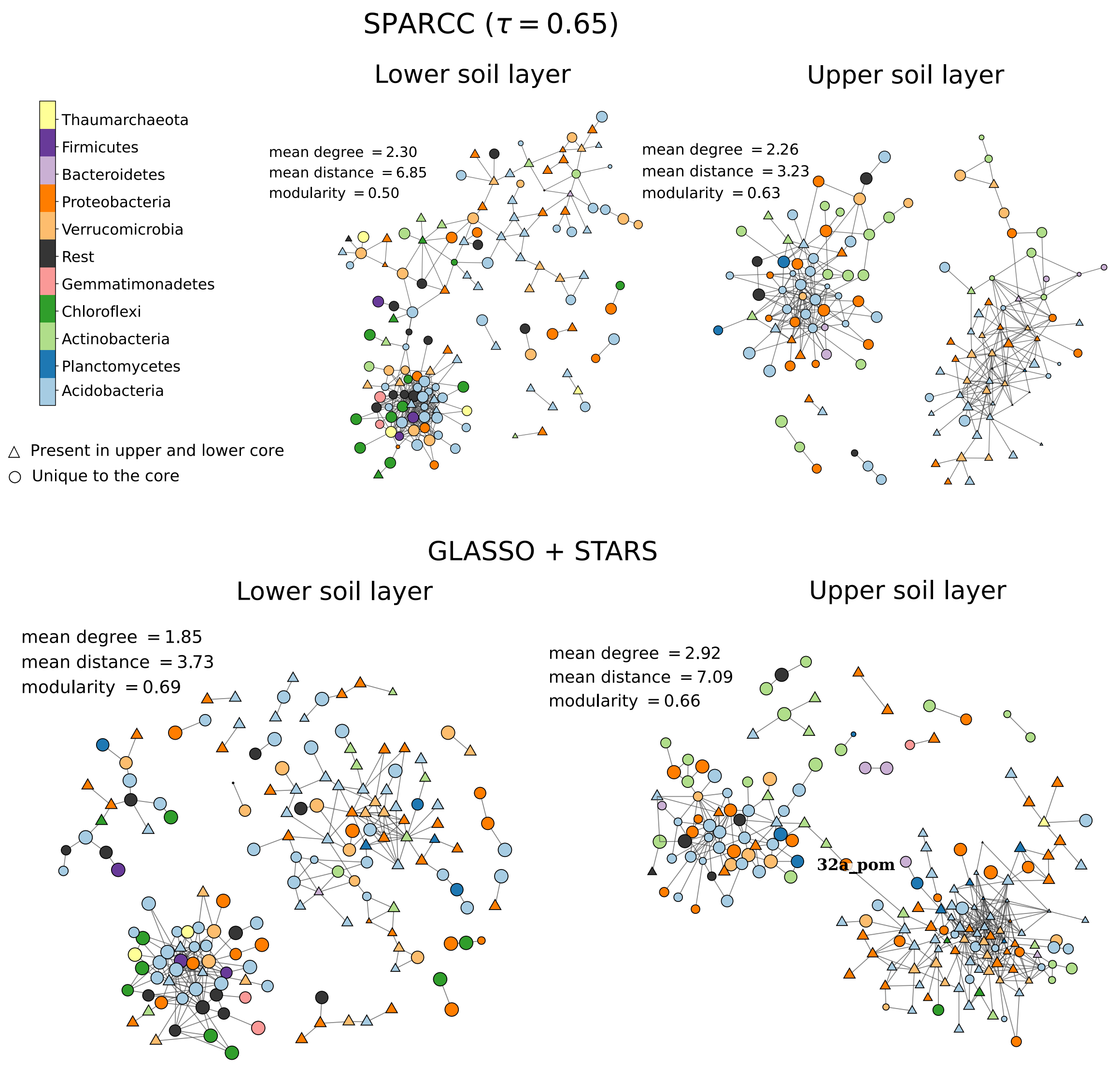


FIG. S8: Networks of positive associations for microorganisms from the upper and lower soil layer detected by: (upper panel) SparCC (lower panel) SPIEC-EASI-like with phylogenetic classifications of nodes (complements Fig.4 of the main text).

[1] Callahan, B. J., McMurdie, P. J., Rosen, M. J., Han, A. W., Johnson, A. J. A., and Holmes, S. P. (2016). DADA2: High-resolution sample inference from Illumina amplicon data. Nature Methods, 13(7):581–583.

[2] Davis, N. M., Proctor, D. M., Holmes, S. P., Relman, D. A., and Callahan, B. J. (2018). Simple statistical identification and removal of contaminant sequences in marker-gene and metagenomics data. Microbiome, 6(1):226.

[3] Eichorst, S. A., Strasser, F., Woyke, T., Schintlmeister, A., Wagner, M., and Woebken, D. (2015). Advancements in the application of NanoSIMS and Raman microspectroscopy to investigate the activity of microbial cells in soils. FEMS Microbiology Ecology, 91(10).

[4] McMurdie, P. J. and Holmes, S. (2013). phyloseq: An R Package for Reproducible Interactive Analysis and Graphics of Microbiome Census Data. PLOS ONE, 8(4):e61217. Publisher: Public Library of Science.

[5] Pjevac, P., Hausmann, B., Schwarz, J., Kohl, G., Herbold, C. W., Loy, A., and Berry, D. (2021). An Economical and Flexible Dual Barcoding, Two-Step PCR Approach for Highly Multiplexed Amplicon Sequencing. Frontiers in Microbiology, 0.
